# A morphological transformation in respiratory syncytial virus leads to enhanced complement activation

**DOI:** 10.1101/2021.05.06.442421

**Authors:** Jessica P. Kuppan, Margaret D. Mitrovich, Michael D. Vahey

**Author notes:** These authors contributed equally to this work (Jessica P. Kuppan, Margaret D. Mitrovich).

## Abstract

The complement system is a critical host defense against infection, playing a protective role that can also enhance disease if misregulated. Although many consequences of complement activation during viral infection are well-established, specific mechanisms that contribute to activation by different human viruses remain elusive. Here, we investigate complement activation by human respiratory syncytial virus (RSV), a respiratory pathogen that causes severe disease in infants, the immunocompromised, and the elderly. Using a strain of RSV harboring tags on the surface glycoproteins F and G, we were able to monitor opsonization of single RSV particles with monoclonal antibodies and complement components using fluorescence microscopy. These experiments revealed an antigenic hierarchy in complement activation, where antibodies that bind towards the apex of F in either the pre- or postfusion conformation are able to activate complement whereas other antibodies are not. Additionally, among antibodies that were able to activate complement, we observed preferential targeting of a subset of particles with globular morphology, in contrast to the more prevalent viral filaments. We found that enhanced complement activation on these particles arises from changes in surface curvature that occur when the viral matrix detaches from the surrounding membrane. This transformation occurs naturally over time under mild conditions, and correlates with the accumulation of postfusion F on the viral surface. Collectively, these results identify antigenic and biophysical characteristics of virus particles that contribute to the formation of immune complexes, and suggest models for how these factors may shape disease severity and adaptive immune responses to RSV.

## Introduction

The complement system is a network of proteins that play a vital role in the innate and adaptive immune responses to pathogens including viruses^1–3^, bacteria^4,5^, and parasites^6–8^. Activation of the complement system proceeds through three principal routes: the classical, lectin, and alternative pathways (reviewed in ref. 9). These pathways differ in their mechanisms of activation: by opsonizing antibodies (classical pathway), by pathogen-specific carbohydrates (lectin pathway), or through continual low levels of attachment to surfaces (alternative pathway). Following activation, each pathway converges on C3, the central component of the complement cascade. C3 that has been cleaved by proteases or that has spontaneously hydrolyzed can covalently attach to activating surfaces via a reactive thioester. C3 attachment to surfaces is self-amplifying, producing new C3 convertases that further drive opsonization. C3 attachment also contributes to the terminal arm of the complement cascade, eventually leading to the assembly of a membrane attack complex that can neutralize membrane-bound targets through the formation of lytic pores.

In addition to its role in the neutralization of pathogens and infected cells, C3 also plays a central role in immune signaling. Activation of complement results in the cleavage of C3 into two fragments, C3a and C3b. C3b attaches to pathogen surfaces, where its multiple degradation products interact with a variety of immune receptors. These interactions contribute to antigen transport to and within lymphoid organs^10,11^; antigen presentation by follicular dendritic cells^12^; activation of B cells^13,14^; T cell priming^2^; and the clearance of immune complexes by phagocytic cells^15^ or erythrocytes^16^. Additionally, C3a that is produced during complement activation is a potent anaphylatoxin, increasing inflammation and recruiting immune cells to sites of infection^17,18^. The diversity of interactions between immune cells and C3 highlights its central protective role bridging innate and adaptive immunity, as well as the potential dangers associated with misregulation of complement^19,20^. Although the disparate contributions complement makes to health and disease are well-established, the mechanisms by which different human viruses activate or evade complement are less well understood. Understanding the factors that contribute to this process could help aid in the development of more effective vaccines and improve understanding of disease pathogenesis.

Here, we set out to investigate complement activation by the human pathogen respiratory syncytial virus (RSV). Activation of complement during RSV infection has been linked to both protection^21,22^ and pathogenesis^17,23^, but the mechanisms that drive complement activation by RSV remain unclear. RSV is an enveloped, negative-sense single-stranded RNA virus of the family *Pneumoviridae* that causes severe infection among infants, the immunocompromised, and the elderly. The RSV genome encodes three membrane proteins – the fusion protein (F), the attachment glycoprotein (G), and the short hydrophobic protein (SH) - which are expressed on the surface of infected cells and packaged to varying degrees into shed virus particles. Among these surface proteins, F and G serve as the primary targets of the adaptive immune response, as well as the leading candidates for vaccine development and prophylaxis^24–27^. The fusion protein, F, mediates the merger between viral and cellular membranes during RSV entry^28^, and has recently been reported to induce outside-in signaling via IGF-1R to facilitate this process^29^. The glycoprotein, G, mediates attachment through the chemokine receptor CX3CR1^30–32^ and, when expressed in soluble form, helps antagonize immune responses^22^. While antibodies against F can provide potent protection both *in vitro* and *in vivo*, the conformational rearrangements this protein undergoes have historically presented challenges in vaccine design^33^. Although antibodies against the prefusion conformation of F (pre-F) are frequently capable of blocking viral entry, antibodies that bind to the postfusion conformation of F (post-F) often fail to do so^34^. Numerous recent breakthroughs in protein design are helping to overcome this challenge through the development of stabilized pre-F antigens as vaccine candidates^35–37^. However, in the context of natural infection, both pre- and post-F conformations occur, and high antibody titers against post-F have been associated with enhanced disease severity and increased activation of complement in the lungs^23,38^.

Understanding how antibodies and RSV antigens contribute to complement activation could provide new insights into vaccine development and mechanisms of RSV pathogenesis. To investigate how RSV-specific antibodies contribute to activation of complement, we developed a fluorescence imaging-based approach to simultaneously quantify antibody binding, the abundance of viral antigens, and the deposition of complement proteins on RSV at the single-virus level. These experiments identify an antigenic hierarchy for complement activation that is dictated by accessibility of the antibody Fc for binding by C1, the initiator of the classical pathway. We also identify a role for the complement defense protein CD55 (DAF), which is packaged into virus particles and increases complement activation thresholds and decreases C3 deposition. Finally, we identify biophysical features of individual RSV particles within heterogeneous populations that enhance their tendency to activate complement. In particular, we find that the detachment of the RSV matrix from the viral envelope – a transition that can be induced by physical perturbations but also occurs under normal physiological conditions *in vitro* – significantly enhances complement activation by a range of antibodies targeting either pre-F, post-F, or both conformations of F. Collectively, these results identify a constellation of mechanisms that contribute to complement activation and immune complex formation by RSV, and inform models for how complement may shape disease severity and the adaptive immune response to infection.

## Results

### Complement activation by F-specific antibodies varies by antigenic site

To identify determinants of complement activation by RSV, we developed a fluorescence assay to quantify antibody binding and opsonization with complement C3 across populations of individual virus particles. Following our previous work using influenza A virus^39^, we engineered a strain of RSV A that was amenable to fluorescence imaging of infected cells and shed virus particles (Supporting information, Figure S1A). To track viral infection, we inserted a fluorescent reporter (mTagBFP2) downstream of the NS1 stop codon but upstream of the gene-end sequence using an internal ribosome entry site. Separately, we inserted a ybbR-tag at the C-terminus of G and a pentaglycine tag immediately following the signal sequence in F. Sfp synthase^40^ can be used to conjugate CoA-based fluorescent probes to the tag on G, while the tag on F becomes exposed following cleavage of the signal sequence, creating a suitable substrate for the enzyme Sortase A (SrtA) to conjugate small peptide-based fluorophores^41^. These modifications allow us to visualize RSV particles and measure densities of antigen (*i*.*e*. F or G) on the viral surface independent of conformational changes in F (*i*.*e*. prefusion vs. postfusion) and while preserving antigenic sites. Viruses labeled in this way retain levels of infectivity matching unlabeled controls (Figure S1B), demonstrating that the attachment of fluorophores is non-disruptive.

Using this system, we sought to characterize activation of the classical pathway by RSV particles. We enzymatically labeled F on the surface of RSV particles collected from A549 cells and immobilized these viruses onto pegylated coverslips functionalized with the anti-G antibody 3D3^25^. This allowed us to quantify opsonization following incubation of fluorescent viruses with normal human serum supplemented with defined fluorescent mAbs and labeled C3 (Figure 1A). Initial tests using normal human serum resulted in robust C3 deposition in the absence of supplemental mAbs (Figure S1C, left column). Incubating virus with this serum for 30 minutes at 4°C also inhibited binding of a high-affinity pre-F-specific mAb (5C4) ∼2-fold and a post-F-specific mAb (ADI-14359) by ∼10-fold, suggesting the presence of competing polyclonal IgG/IgM within the serum (Figure S1D). In contrast, IgG/IgM-depleted normal human serum showed little C3 deposition in the absence of supplemental mAbs, but robust opsonization with C3 when F-specific mAbs were added (Figure S1C, right column). IgG/IgM-depleted serum therefore provides a means of determining the ability of individual mAbs to activate complement, allowing us to identify antibody features that are predictive of potency.

**Figure 1:**
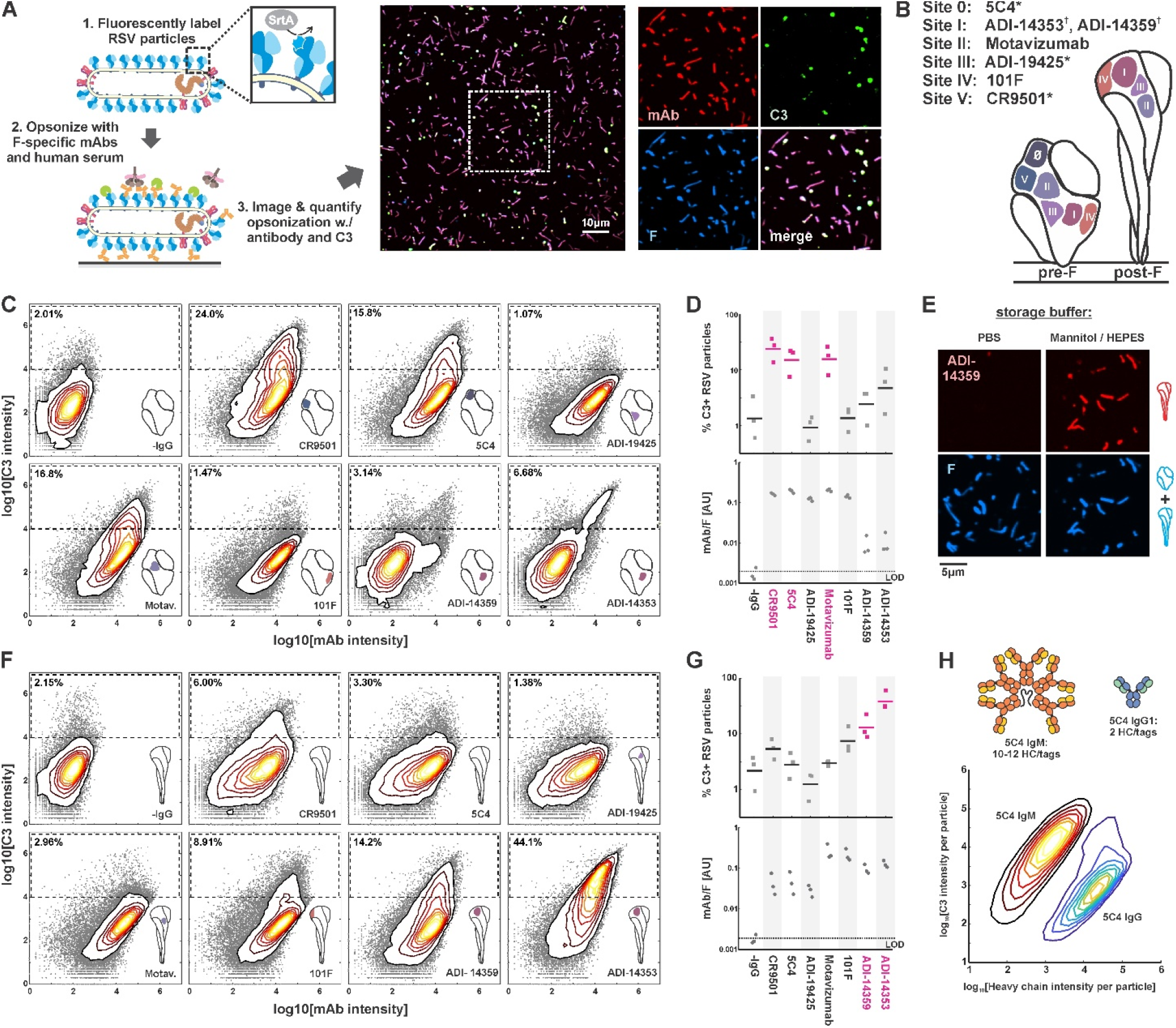
Complement activation and C3 deposition vary across antigenic sites of RSV F. (A) A fluorescence-based approach to measuring opsonization of RSV particles with mAbs and C3. RSV particles with site-specifically labeled F are immobilized on coverslips and incubated with normal human serum (IgG/IgM-depleted), specific mAbs, and fluorescent C3 prior to imaging. *Right*: RSV particles opsonized with mAb (CR9501 IgG1 in the image shown) and C3. (B) Antibodies used in this study and their antigenic sites. A ‘*’ denote mAbs specific to prefusion F while a ‘^†^’ denotes mAbs specific to postfusion F. (C) Distributions of integrated antibody and C3 intensities on opsonized virus particles with predominantly prefusion F. Gray points indicate data for individual virus particles. Dashed regions indicate criteria for C3-postiive particles. Data is combined from three biological replicates. (D) *Top*: Data from *C*, plotted as percentage of C3-positive RSV particles, defined by integrated intensities >10^4^. Points show results from three biological replicates with the mean across the replicates shown as a line. Antibodies considered activators of the classical pathway (>10% C3-positive particles) are shown in magenta. *Bottom:* Plot of average mAb:F intensity per RSV particle across the same antibodies and replicates as in the plots above. (E) Conversion of pre-F to post-F on RSV filaments via ∼24h incubation in buffer with low ionic strength but balanced osmolarity. Post-F is detected using the site I-directed mAb ADI-14359. Images are displayed at matching contrast levels. (F) and (G): Results corresponding to *C* and *D* but for RSV particles containing predominantly post-F. (H) Comparison of C3 deposition by IgM and IgG1 antibodies. *Top*: schematic of antibodies based on 5C4. The antibodies contain matching light chains and VH domains coupled to the human heavy chain μ (for IgM), or a human IgG1 Fc. IgM is additionally co-expressed with a J chain plasmid. All heavy chains contain a c-terminal ybbR tag for site-specific conjugation of a fluorophore for quantification of bound antibody. *Bottom*: contour plots showing distributions of C3 and IgM/IgG1 heavy chains per RSV particle, based on fluorescence intensities. The IgM distribution is determined from 317594 RSV particles; the IgG distribution is determined from 126372 RSV particles.

We expressed and purified a panel of human IgG1 mAbs targeting each of the known antigenic sites of RSV F^26^ (Figure 1B). Each mAb was modified to contain a ybbR-tag^40^ at the C-terminus of the heavy chain, to permit quantitative site-specific labeling and the ability to measure the amounts of antibody bound to individual virus particles. By using mAbs at concentrations that saturate binding, we were able to decouple binding affinity from the intrinsic capacity of a given mAb to activate complement once bound. Under conditions that saturate binding, these mAbs vary markedly in their ability to activate complement. Among the pre-F-binding mAbs tested, 5C4^24^ (site 0), CR9501^42^ (site V), and Motavizumab^43^ (site II) showed the greatest potency. In contrast, ADI-19425^34^ (site III) and 101F^44^ (site IV) showed only background levels of C3 deposition (Figure 1C & D). Thus, although each pre-F-binding mAb achieved similar levels of binding to RSV F (Figure 1D, lower plot), only three out of five activated complement above background levels.

Several F-specific antibodies bind to the postfusion conformation of F, either exclusively or in addition to the prefusion conformation. This includes several antibodies in our panel: 101F and Motavizumab (which bind to both pre- and post-F) as well as ADI-14359 and ADI-14353 (which bind specifically to post-F)^34^. To compare C3 deposition driven by these antibodies upon binding to post-F, we first incubated viruses bound to coverslips for 24h in buffer with low ionic strength (300mM mannitol, 10mM HEPES pH 7.2), based on the previous finding that pre-F spontaneously triggers to post-F in buffers with low salt concentrations^45,46^. This treatment increased binding of the post-F specific antibodies ADI-14359 and ADI-14353 approximately 300-fold without obvious changes in the morphology of virus particles (Figure 1E). Conversion of RSV particles to a predominantly post-F form increased C3 deposition from post-F specific mAbs ADI-14359 and ADI-14353 to levels comparable to or greater than the most potent pre-F specific antibodies in the context of pre-F antigens (Figure 1F & G). For the conformation-independent mAbs 101F and Motavizumab, C3 deposition following pre-to-post-F conversion followed different trends, increasing approximately six-fold for 101F but decreased by a similar ratio for Motavizumab, despite similar levels of antibody binding (Figure 1F & G).

The classical pathway is activated when the C1 initiation complex binds to IgM or IgG that has assembled on activating surfaces. While activation via IgG requires the assembly of a hexameric complex of antibody Fc regions^47–49^, IgM is pre-assembled as an activating platform, with C1 binding sites exposed only upon engagement with surface antigen^50^. As a comparison with our IgG antibodies, we tested activation of the classical pathway by recombinant IgM with VH and VL domains from 5C4. As expected, this high-affinity IgM was considerably more potent than its IgG1 counterpart, leading to >100-fold more C3 deposition per bound heavy chain than the corresponding IgG1 mAb (Figure 1H). Given the importance of establishing a platform for C1 binding in activation of the classical pathway, we sought to identify how the IgG antibodies in our panel may differ in this regard.

A consistent trend among the IgG1 antibodies that most efficiently activate complement deposition is the angle with which they bind to either pre- or post-F. 101F and Motavizumab, antibodies that bind to both conformations but with alternating Fc orientations, illustrate this effect. For both antibodies, C3 deposition increases approximately six-fold when the Fc is oriented away from the viral membrane as opposed to towards it. Projection of the Fc region further above the viral membrane is common to all of the activating antibodies we tested (Figure S2A), and potentially increases accessibility for binding by C1. Moreover, positioning the Fc regions on a plane above the surrounding canopy of F could also facilitate Fc hexamer formation^47^ by avoiding steric hindrance from neighboring proteins in the viral membrane. Consistent with this model, we observe more binding by C1 (as detected by an anti-C1q antibody) to mAbs that bind to pre-F with their Fcs projected outward (CR9501, 5C4, Motavizumab) compared to those where the Fc lies within or below the plane of F trimers in the viral membrane (ADI-19425, 101F) (Figure S2B & C). Of note, we still observe C1 binding for mAbs that do not drive C3 deposition (*e*.*g*. ADI-19425), suggesting that attachment of C1 to these mAbs may be less likely to produce an active C1 complex. This could occur if a high density of antibodies permitted C1 to attach, but assembly of the activating hexamer were occluded by F, G, or other proteins present at high densities on the viral surface.

### CD55 is packaged into RSV particles and modulates sensitivity to complement deposition

Complement defense proteins anchored in the membranes of host cells can be packaged into enveloped viruses during assembly^51–53^, where they may function to restrict different stages of the complement cascade. To determine if host complement defense proteins restrict opsonization of RSV particles with C3, we focused on the roles of CD46 and CD55. Both proteins are abundantly expressed on A549 cells and function to limit the amplification step of the complement cascade by restricting the formation or stability of new C3 convertases^54^. Using fluorescent Fab fragments or antibodies against CD46 and CD55, we were able to detect both on the surface of RSV particles collected from A549 cells. We proceeded to construct two polyclonal A549 knockout cell lines where one gene or the other was deleted via CRISPR/Cas9. Following knockout and cell sorting, we were no longer able to detect the targeted proteins in cells or shed viruses, verifying the specificity of the antibodies and confirming successful knockout (Figure 2A&B). Using these cells lines, we proceeded to compare C3 deposition in the presence or absence of CD55 or CD46. Virus released from CD55 KO cells showed increased sensitivity to C3 deposition across antibodies specific to site 0, II, and V, with percentages of opsonized particles increasing ∼2-3 fold relative to virus produced by wildtype cells (Figure 2C). Conversely, virus released from CD46 KO cells did not show significant differences in C3 deposition relative to wildtype cells using the site-V-specific mAb CR9501 (Figure 2D). Comparing activation by CR9501 across a range of antibody dilutions suggests that deletion of CD55 increases opsonization to an extent comparable to a 4-fold increase in antibody concentration (Figure 2E&F). While additional complement defense proteins may play critical roles in other aspects of RSV infection, these results show that CD55 plays an outsized role in modulating sensitivity to opsonization with C3.

**Figure 2:**
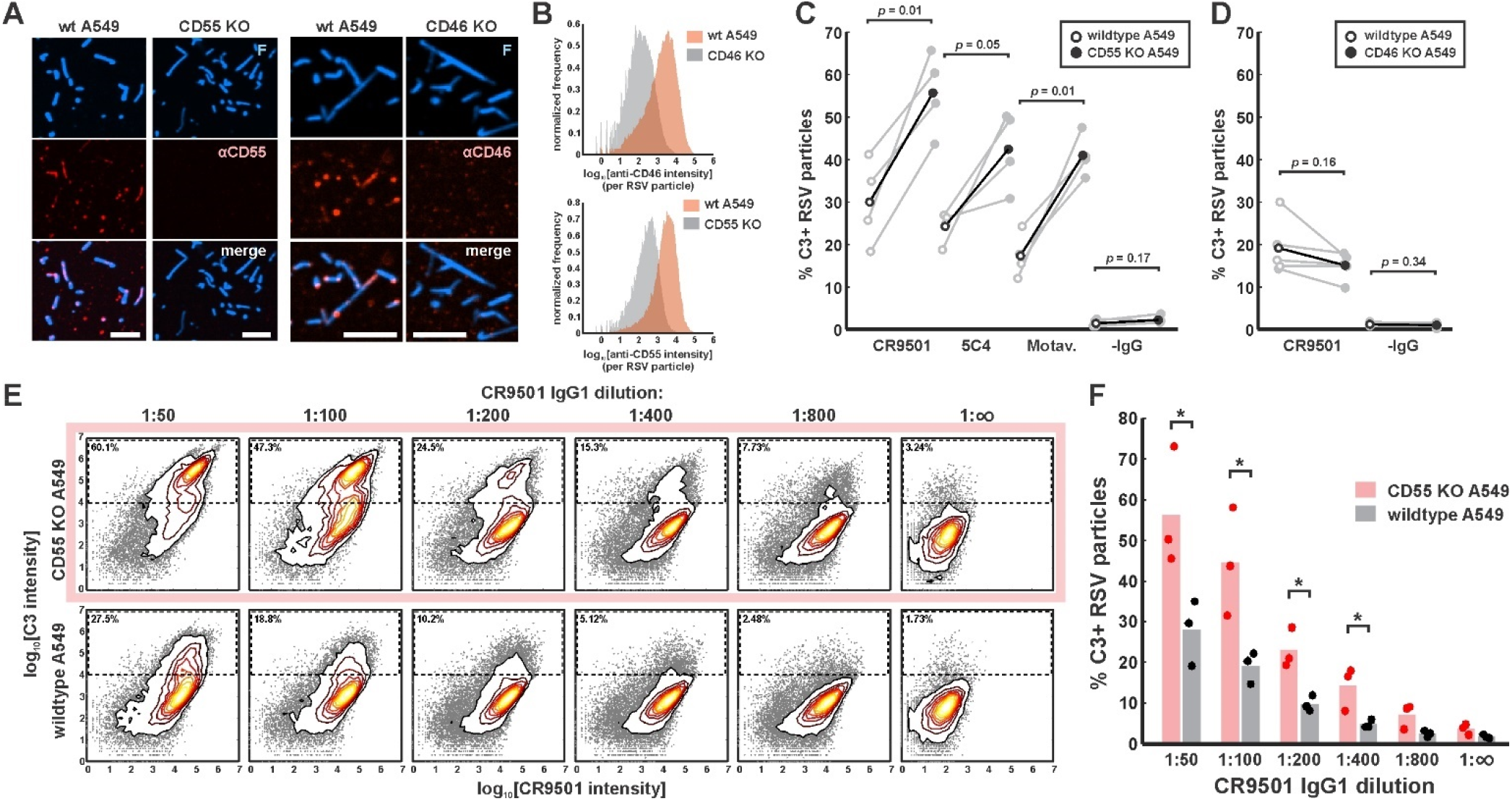
CD55 is packaged into RSV particles and increases thresholds for C3 opsonization. (A) Images of RSV particles with enzymatically-labeled F and CD55 or CD46 labeled via fluorescent antibodies. Panels show representative images of virus released from wildtype (wt) A549 cells and CD55 or CD46 knockout cells, displayed at matching contrast levels. Scale bar = 5μm. (B) Histograms showing distributions of antibody intensities per RSV particle for wildtype and knockout cell lines. (C) Comparison of C3 deposition for three different F-specific antibodies and a negative control on viruses raised in wildtype cells (open circles) or CD55 knockout cells (closed circles). Connecting lines show data from paired biological replicates; black lines show mean values. *P*-values are determined using a paired-sample t-test. (D) Similar plot as in *C*, but for data obtained from CD46 knockout cells. (E) Distributions of antibody and C3 intensities for serial dilutions of CR9501 IgG1. Panels in the top row show results for viruses from CD55 KO cells, while panels from the bottom row show results for viruses from wildtype cells. The region indicated by the dashed line represents the threshold for C3-positive particles. Percentages in the upper left corners indicate the percentage of C3-postiive particles. Data is combined from three biological replicates. (F) Data from *E* plotted as percentage C3-positive particles per condition. * indicates *p*<0.05 calculated using a two-sample t-test.

### Globular particles containing postfusion F serve as dominant targets of complement activation by both pre-F and post-F specific antibodies

Although the efficiency of opsonization with C3 varies depending on the activating antibody and on the presence or absence of host complement defense proteins, we observed a consistent pattern across experimental conditions, where particles opsonized with C3 had more globular morphology than those with low or undetectable levels of C3, which tended to be more filamentous (Figure 3A & B). To investigate this further, we characterized particle morphology and F conformation by simultaneously labeling RSV particles with the prefusion-specific antibody 5C4 and the postfusion-specific antibody ADI-14359. This revealed that filamentous particles contained almost exclusively pre-F while globular particles were frequently enriched in post-F (Figure 3C), consistent with prior characterization using electron microscopy^55^. Comparing newly-released virus with virus incubated in cell culture media at 37°C for 24 hours revealed that the proportion of post-F-containing particles increased following the 24h incubation, while the proportion of filamentous particles (defined as those >1um in size with an aspect ratio >2) decreased (Figure 3D-F). Direct labeling of RSV-infected cells with 5C4 and ADI-14359 revealed mixtures of pre- and post-F particles on the surfaces of both infected A549 cells as well as on neighboring uninfected cells, confirming that the occurrence of both particle types is not limited to circumstances where the virus is being collected or handled (Figure S3). Collectively, these results suggest that RSV particles transform spontaneously over time under mild conditions into a globular state enriched in post-F.

**Figure 3:**
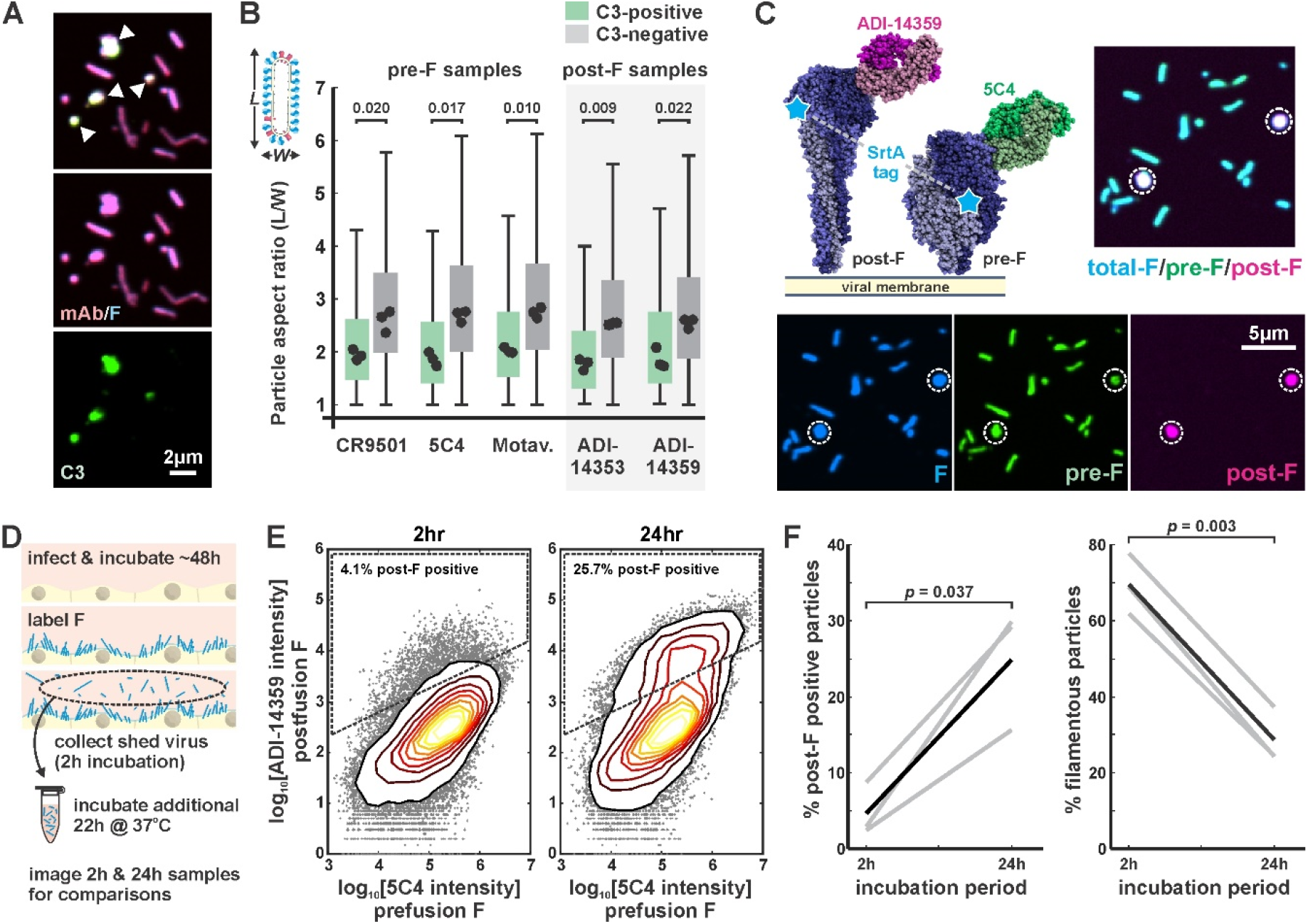
RSV particles are morphologically and antigenically heterogeneous. (A) RSV particles opsonized with antibody (CR9501 IgG1) and C3. White arrows in merged panel indicate globular particles with high levels of C3. (B) Comparison of particle aspect ratios across RSV particles that do (green boxes) or do not (gray boxes) show opsonization with C3. Boxes indicate 25^th^-75^th^ percentiles and points show median values for three biological replicates. Analysis is limited to particles with an area larger than 100 pixels, where morphology can be determined from diffraction-limited images. *P*-values are determined based on median values using a paired sample t-test. (C) Three-color labeling strategy to detect total F (via enzymatic labeling with SrtA), post-F (via the post-F specific mAb ADI-14359) and pre-F (via the pre-F specific mAb 5C4). Fluorescence images show RSV particles labeled to indicate pre-F, post-F, and total F on the virion surface. (D) Experimental approach to determine effects of aging on RSV particles. (E) Distributions of pre-F and post-F intensities for virus aged 2h at 37°C (left) or 24hr at 37°C (right). Data is combined from three biological replicates. Region inside the dashed lines define criteria for post-F positive particles. (F) Percentage of post-F positive particles (left) and filamentous particles (right) after 2h and 24h aging. Post-F-positive particles are defined by those within the dashed lines in *E*. Filamentous particles are defined as those with length >1μm and aspect ratio (*L/W*) > 2. Gray lines show results from paired biological replicates; black lines show average values. *P*-values are determined using a paired sample t-test.

The bias in C3 opsonization towards globular particles could reflect a higher intrinsic sensitivity to complement activation in these particles as compared to viral filaments, or it could indicate that particle morphology is altered upon deposition of C3 or other complement proteins. To distinguish between these two possibilities, we compared C3 deposition on newly-shed virus (collected over a 2-hour window) to “aged” virus (incubated at 37°C an additional 22 hours following release from cells). If globular morphology predisposes particles to C3 deposition, we would expect to see more C3 deposition in the particles aged an additional 22 hours. Consistent with this prediction, we observed that virus collected at 2h (retaining a high proportion of filaments) was less sensitive to opsonization than virus aged an additional 22h at 37°C (retaining a lower proportion of filaments) (Figure 4A & B), suggesting that particles with globular morphology and/or higher levels of post-F have lower thresholds for complement activation. This trend is conserved even for the pre-F-specific mAb CR9501, an unexpected result given that samples incubated at 37°C show ∼20% loss in pre-F as it converts to post-F (Figure 3F, Figure 4B).

**Figure 4:**
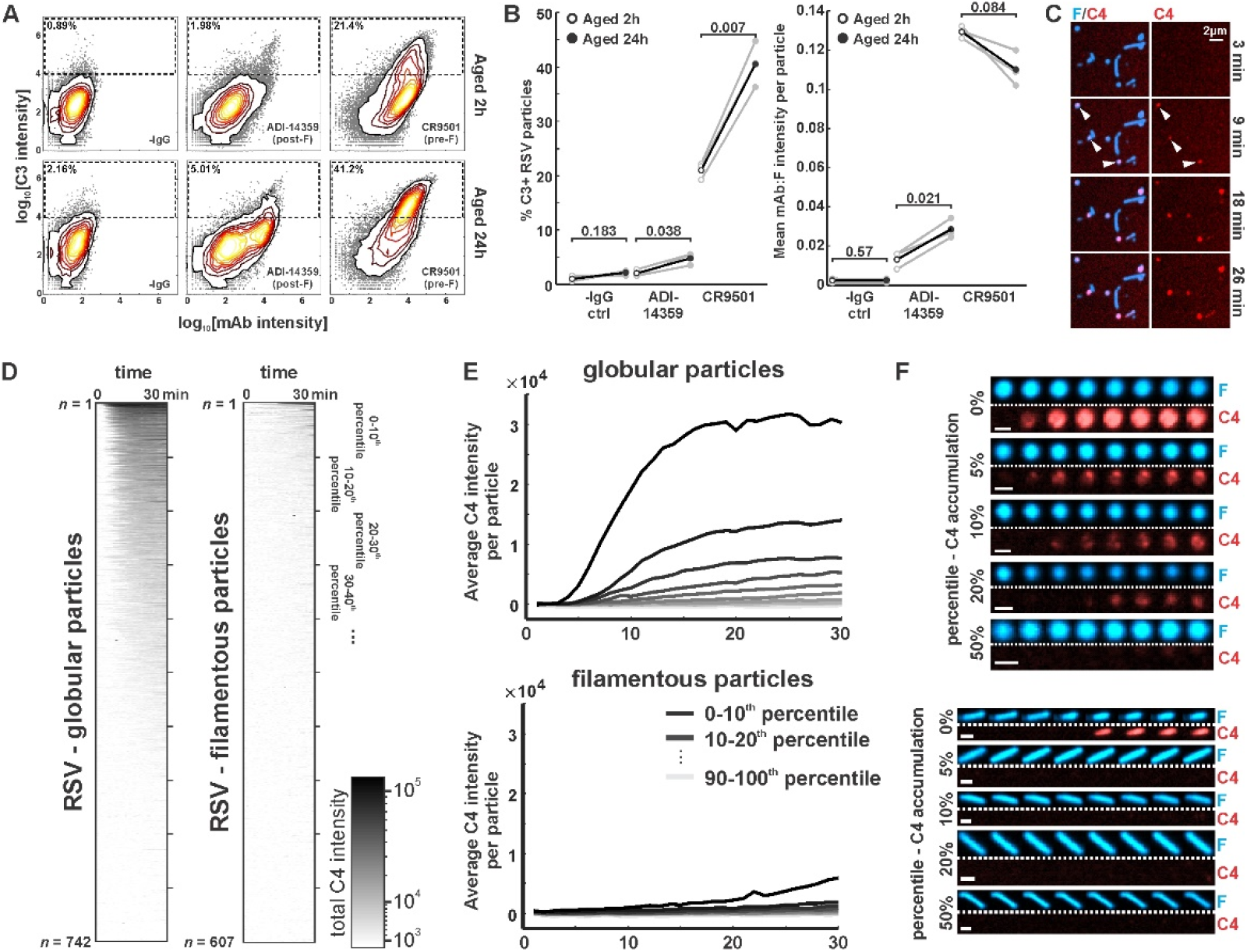
Globular particles are preferentially and rapidly opsonized with C3 and C4. (A) Distributions of C3 and mAb intensities per RSV particle in the absence of mAb (‘-IgG’) or in the presence of ADI-14359 or CR9501. The top row shows results for particles aged for a total duration of 2h at 37°C (i.e. during budding from A549 cells); the bottom row shows particles aged for a total duration of 24h at 37°C following collection at 2h. Particles within the dashed rectangles indicate those that are C3-positive, and the percentage of these particles is indicated in the upper left. Distributions are combined from three biological replicates. (B) *Left*: percentage of C3-positive particles following aging at 37°C for 2h (open circles) or 24h (filled circles). Black markers give average values for three biological replicates; individual replicates are shown in gray. *Right*: mean mAb/F intensities per particle for the same datasets plotted to the left. *P*-values determined using a paired-sample t-test. (C) Time series of C4 accumulation on RSV particles. White arrows indicate first detectable accumulation of C4. (D) C4 accumulation on RSV particles over time, categorized by particle morphology. Data to the left shows C4 accumulation on 742 globular particles; data on the right show C4 accumulation for 607 filamentous particles. In both plots, rows show C4 data for individual particles from 0 to 30 minutes, sorted from top to bottom according to final C4 accumulation. (E) Data from *D*, plotted by grouping particles within percentile intervals from 0-10%, 10-20% etc. and averaging C4 accumulation in each group. (F) Sample images of globular (top images) and filamentous (bottom images) RSV particles. Displayed images are sampled at 4 minute intervals beginning from 2 minutes after the addition of serum and complement components (scale bar = 1μm).

To further confirm that differences in particle morphology precede differences in opsonization, we performed time-resolved experiments to determine the kinetics of complement deposition. For these experiments, we used CR9501 mAb as the activating antibody and tracked deposition of C4 on virus particles rather than C3. The lower concentrations of C4 in serum as compared to C3 allowed us to visualize deposition on particles without the high background signal from protein in solution that arises when using C3. Additionally, deposition of C4b is immediately downstream of activation of the C1 complex in the classical pathway, providing rapid detection of complement activation. Consistent with endpoint measurements of C3 deposition, C4 accumulates first on globular particles, appearing within ∼10-15 minutes of incubation with complement components (Figure 4C). Analysis of C4 deposition across *n* = 742 globular and *n* = 607 filamentous particles revealed substantial differences in patterns of opsonization; ∼50% of globular particles showed detectable accumulation of C4 within 30 minutes, compared to <10% of filamentous particles (Figure 4D-F). Moreover, opsonization with C4 proceeds with different kinetics, occurring ∼5-fold faster in globular particles than in filamentous ones (Figure 4E&F). These results demonstrate that sensitivity to antibody-dependent complement deposition correlates with RSV particle morphology.

### Detachment of the viral matrix increases complement activation by decreasing membrane curvature

The globular RSV particles we observe resemble prior observations from electron microscopy, where the matrix protein appears to have dissociated from the membrane, resulting in more rounded morphology and less ordered distributions of surface proteins as compared to viral filaments budding from infected cells^55,56^. Although this effect has been attributed to damage during sample preparation, we reasoned that aging may result in the same morphological transformation, and that a similar effect could be achieved in a controlled fashion through osmotic swelling. Treatment of RSV particles with a low osmolarity buffer (10mM HEPES pH 7.2, 2mM CaCl_2_) transformed viral filaments into spherical particles over the course of ∼1 min, with no loss of infectivity (Figure 5A, Figure S4A). To corroborate that this morphological transformation coincides with detachment of the viral matrix, we performed photobleaching experiments on RSV particles with enzymatically labeled F. In the absence of an intact matrix layer, we reasoned that F (which may interact with the matrix protein via its cytoplasmic tail^57,58^) could freely diffuse laterally within the plane of the viral membrane and should therefore show increased recovery after partial photobleaching (Figure S4 B&C). Consistent with this prediction, we observed significantly more recovery in bleached F on post-F enriched globular viruses and on osmotically-swollen viruses compared to viral filaments. The increased recovery on osmotically swollen viruses could be reversed by treatment with 0.5% PFA while preserving the particles’ spherical geometry (Figure S4C). These results suggest that the morphological transformation we observe as RSV particles age or are subjected to physical perturbations is driven by the detachment of the matrix from the viral membrane.

**Figure 5:**
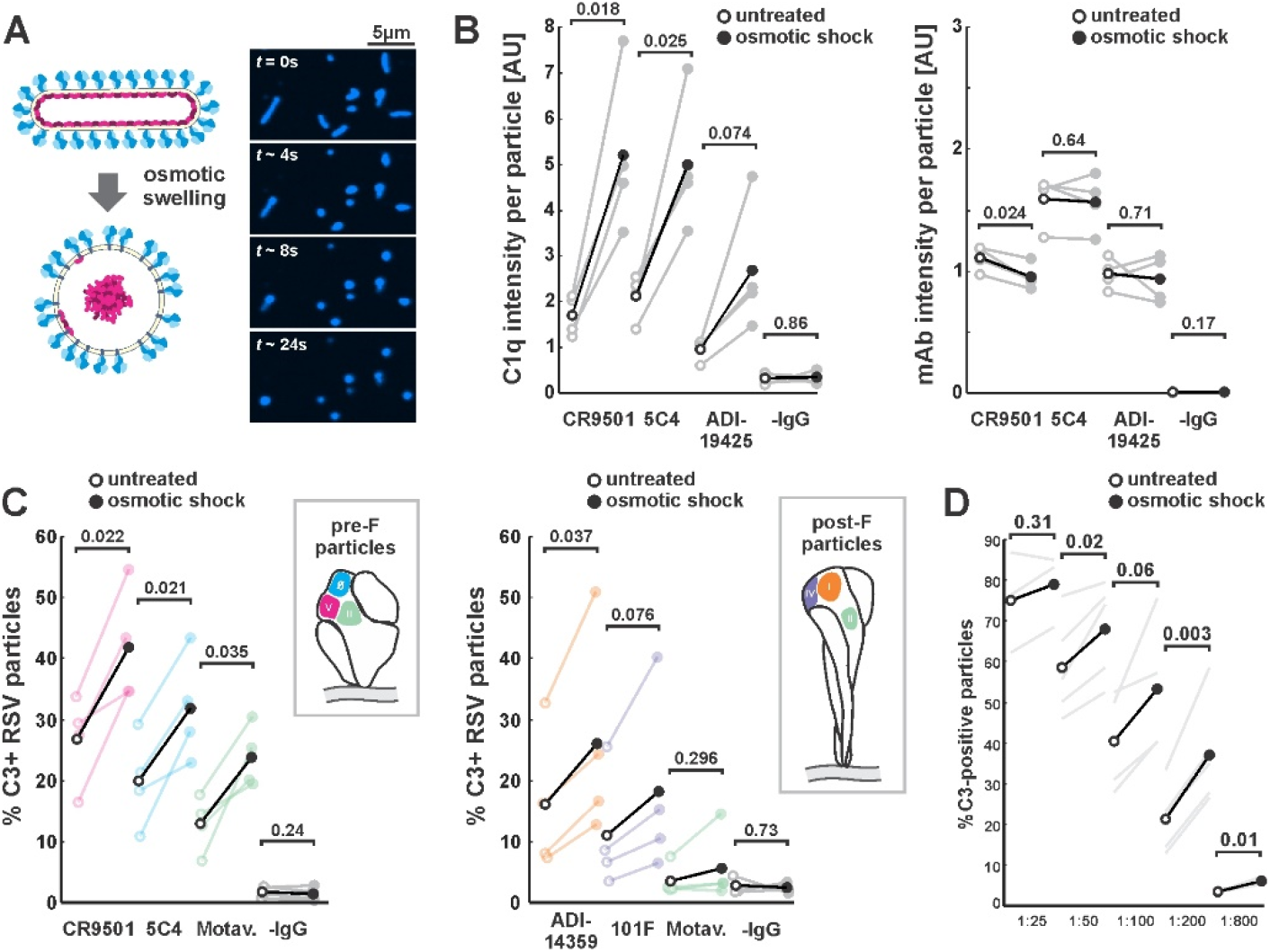
Membrane detachment from the viral matrix increases complement activation. (A) Process for detaching the RSV matrix from the viral membrane using osmotic swelling. Fluorescence images show details of the transformation, with *t* = 0s approximately corresponding to the addition of low osmolarity buffer. (B) *Left*: C1q binding per RSV particle, with (filled circles) and without (open circles) osmotic swelling. Black markers indicate averages across four biological replicates. Individual replicates are shown in gray, with connecting lines indicating paired replicates. C1q intensity is measured using a C1qA-specific antibody conjugated to Alexa Fluor 488. *Right*: average mAb intensity per particle for the same experiments plotted to the left. *P*-values determined using a paired-sample t-test. (C) Percent C3-positive RSV particles with (filled circles) and without (open circles) osmotic swelling. Plots to the left are for predominantly pre-F particles; plots to the right are for predominantly post-F particles. Black markers indicate averages of three biological replicates (color-coded according to the mAb’s antigenic site). *P*-values determined using a paired-sample t-test. (D) Percent C3-positive RSV particles with (filled circles) and without (open circles) osmotic swelling across different dilutions of 5C4 IgM. *P*-values determined using a paired-sample t-test.

We next sought to determine the effects of this controlled transformation on complement activation. We found that osmotic swelling led to increased C1 binding in the presence of F-specific mAbs (CR9501, 5C4, ADI-19425), but not in their absence (Figure 5B, left), indicating that the effect is not a non-specific consequence of swelling. Furthermore, increased C1 binding did not result from increased mAb binding, which remains constant or decreases slightly upon swelling (Figure 5B, right). Similar to the effects of aging on virus particles (Figure 4), C3 opsonization also increases upon osmotic swelling, an effect that is conserved across mAbs targeting a range of antigenic sites on pre-F and/or post-F (Figure 5C). Recombinant IgM antibody with VH and VL domains from 5C4 shows a similar effect, with C3 deposition increasing by ∼50% following osmotic swelling across a range of IgM dilutions (Figure 5D). Collectively, these results demonstrate that detachment of the viral matrix increases sensitivity to complement activation by both IgG and IgM antibodies, and that increased sensitivity is likely due in part to increased attachment of C1 to the viral surface.

The morphological transformation in RSV particles is accompanied by two notable biophysical changes: increased mobility of F in the membrane and decreased membrane curvature. While either could potentially contribute to enhanced complement activation, we observed a similar fold-increase in C3 deposition on both fixed (0.5% PFA) and unfixed viruses in the spherical state vs. the filamentous state (Figure S4D), suggesting the antigen mobility alone does not account for preferential opsonization of globular RSV particles. We therefore sought to determine if membrane curvature could be a determining factor. Curvature has previously been implicated in complement activation on antigen/antibody-coated beads and peptidoglycan nanoparticles^59–61^. However, these studies have reached different conclusions regarding the effects of curvature, and it remains unclear how these results would generalize to enveloped viruses, where antigens are oriented and may be spaced semi-regularly^62–64^.

The transformation from filament to sphere results in decreased membrane curvature to an extent that varies depending on the size of the initial filament. While the change in curvature is negligible for particles whose length and diameter are similar (∼100nm), curvature will decrease ∼2-fold for a particle 1μm in length, and ∼5-fold for particles 10μm in length (Figure S5A & B). If complement activation by IgG is sensitive to curvature, the effects of osmotic swelling on C3 deposition should therefore vary depending on the initial size of the particle. To test this prediction, we compared C3:F ratios between osmotically-swollen (spherical) particles and non-treated (filamentous) particles from the same initial population of RSV. While C3:F ratios are similar for smaller viruses regardless of swelling, C3:F is increased by up to nine-fold upon swelling in larger viruses, indicating a preference for lower curvature in IgG1 activation of complement (Figure S5C). Thus, large viruses are poised to evade complement activation when they emerge from cells as highly-curved filaments, but become substantially more susceptible as they age or their morphology is physically disrupted.

## Discussion

Activation of complement during RSV infection has been linked to both protective and pathogenic effects, but the mechanisms that drive complement activation by RSV remain unclear. We find that a number of factors contribute to activation of the classical pathway by shed RSV particles, including characteristics of the targeting antibody, the packaging of complement defense proteins into the viral membrane, and biophysical characteristics of RSV particles. Among these factors, the dominant contribution comes from the activating antibody. We find an antigenic hierarchy in the ability of different mAbs to activate complement: although all of the IgG1 antibodies we have tested contain the same Fc and are capable of binding to pre- or post-F at similar antibody:F ratios, only a subset are able to activate complement with appreciable efficiency under the conditions of our experiments. A common feature of these activating antibodies is that they are all predicted to project their Fc above the surrounding canopy of F (>15nm above the viral membrane), leading to more efficient binding by C1 (Figure 1, Figure S2). The typical distance between adjacent F trimers in the RSV membrane appears to be ∼10-20nm^56^, a value that is roughly consistent with other filamentous viruses^62,64,65^. Since this spacing is too small to accommodate an activating IgG or IgM platform between F trimers and since F appears to be largely stationary within the membrane of viral filaments (Figure S4B & C), it may be critical to position antibody Fcs in this way so as to avoid steric clashes with adjacent F trimers. A consequence of this scenario is that activation of complement would occur at comparatively large distances from the membrane. While this could potentially limit the proper assembly of a membrane attack complex (where proximity to the membrane may be beneficial^66^), it may be inconsequential to other aspects of activation including the production of C3a, which has been implicated in RSV disease severity^17^.

Perhaps the most unexpected result presented here is the observation that treatments which increase the proportion of post-F in the viral membrane can enhance complement activation by both pre-F and post-F specific antibodies (Figures 4 & 5). Given the importance of C3 opsonization in antigen presentation and B cell activation^14^, we speculate that this may contribute to bias in adaptive immune responses to infection. Previous work has established that infants mount an adaptive immune response focused largely on prefusion F, and that this response shifts over time to recognize post-F as the infants age^34^. Antibodies that bind to pre-F may activate complement disproportionately on post-F containing particles, lowering thresholds for the activation of B cells that engage with these particles. Such an effect could be further enhanced by epitope masking by the pre-F antibodies, which would further reduce the availability of pre-F epitopes for B cell engagement. Although this model is speculative, our results suggest that further investigation of whether or how bias in complement activation contributes to establishing immunodominance hierarchies may be warranted.

Antibody responses specific to postfusion F have been linked to increased disease severity associated with enhanced complement activation^23,38^. Post-F antibodies are poorly neutralizing and thus ineffective at controlling infection, potentially leading to increased viral load. In addition, our results suggest that the high intrinsic capacity of post-F containing particles to activate complement may further enhance the damage caused by post-F specific antibodies. Mechanistically, the increased tendency of post-F enriched particles to activate complement appears to originate from the detachment of the viral matrix, which in turn decreases the mean curvature of the viral surface. Although more work is needed to understand the factors that contribute to matrix detachment and the extent to which this occurs during *in vivo* infection, our observations that these particles accumulate over time under normal cell culture conditions suggest that their presence *in vivo* is feasible, where the physical and chemical environment would be considerably harsher and more complex. Other enveloped viruses, including phylogenetic neighbors of RSV from *Paramyxoviridae* such as measles virus and Newcastle disease virus, have been shown to exhibit a similar pleomorphism arising from matrix disassembly^64,67,68^. More generally, our results suggest that any factor that alters virus morphology – whether genetic or non-genetic - could have biophysical effects on immune signaling.

The methodology presented here - combining site-specific labeling and fluorescence characterization to dissect mechanisms of complement activation by RSV - provides a model that could be generalized to other pathogens. Although investigating complement activation *in vitro* limits our ability to asses direct immunological consequences of activation, it provides the ability to isolate the contributions that different antibodies and distinct particle subsets make towards complement activation. This may be particularly useful for viruses that vary widely in the biophysical characteristics of released progeny, including RSV, HMPV, influenza, and Ebola. Extending this methodology to investigate other viruses and their interactions with immune receptors could help determine if particle heterogeneity contributes to disparate outcomes during immune signaling in other contexts as well.

## Methods

### Creating recombinant RSV for site-specific labelling

We introduced modifications into the RSV genome following the approach of Hotard *et al*.^69^. Briefly, we electroporated a BAC containing the antigenomic cDNA of a chimeric RSV strain A2 with the F protein from Line 19 (obtained through BEI Resources) into the E coli strain SW102^70^. To avoid virus attenuation and fluorescence spectral overlap caused by the mKate2 reporter in front of NS1 in the initial construct, we replaced it with an mTagBFP2 reporter, expressed from an IRES following NS1. From this modified BAC, we proceeded to use a galK cassette with a 5’ homology arm targeting the C-terminal region of G and a 3’ homology arm targeting the N-terminal region of F to insert tags on G and F simultaneously following an initial round of selection on galactose plates. We verified successfully modified BACs by sequencing a PCR-amplified region around the modified site, and we purified the BACs from 250ml cultures using a Nucleobond BAC100 kit. The genomic sequence for the final virus is given in Supporting information.

To rescue recombinant viruses, we transfected 6-well plates of BHK-21 cells with BAC (0.8μg), helper plasmids (codon-optimized L, N, P, and M2-1 at 0.2, 0.4, 0.4, and 0.4μg, respectively), and a vector containing the T7 RNA polymerase (0.2μg) using Lipofectamine 2000. Helper plasmids were obtained from BEI Resources. Following transfection, cells cultured in virus growth media (OptiMEM with 2% FBS and antibiotic-antimycotic) were passaged every 2-3 days at a 1:3 ratio and monitored for signs of infection. To collect virus stocks, we removed media from T75 flasks of infected cells and replaced it with 2ml of PBS supplemented with 2mM EDTA. Cells that detached from the flask were collected, flash-frozen in liquid nitrogen, and pelleted to remove cell debris following a rapid thaw at 37°C. The virus-containing supernatant was then aliquoted and stored at -80°C until needed for experiments. Detailed characterization of the replication of this recombinant strain will be published elsewhere. Viral titers used for MOI calculations were determined by quantifying mTagBFP2-expressing A549 cells infected in 96-well plates as a function of the input volume of virus.

### Preparing viruses

We used viral stocks snap-frozen and stored at -80°C to infect ∼90% confluent A549 cells in 8-chambered coverglass or 96-well plates at MOI ∼1. To enhance the efficiency of infection, virus diluted into virus growth media (final volume of 100ul) was centrifuged onto cells at 1200xg for 10 minutes and returned to the incubator for an additional 50 minutes before washing off the virus-containing media and replacing with fresh virus growth media. Samples used for independent biological replicates were conducted using viruses from independent infections.

Enzymes and probes for site-specific labeling were generated as previously described^39,71^. At 48-60 hours post infection, F on the surface of infected cells was labeled *in situ* using 50μM Sortase A and 100μM fluorescent CLPMTGG substrate. With the exception of photobleaching experiments (where sulfo-Cy3 was used), sulfo-Cy5 maleimide (Lumiprobe, 13380) was conjugated to the N-terminal cysteine of the SrtA peptide to label F. Labeling reactions were prepared in virus growth media supplemented with 5mM CaCl_2_. For experiments with labeled G, the labeling reaction also included 5μM Sfp, 10μM CoA-probe, and 5mM MgCl_2_. After labeling cell surface viral proteins for two hours at room temperature, the labeling reaction was washed four times with fresh media and the cells were returned to the incubator for an additional two hours to allow labeled viruses to detach. Viruses prepared in this way exhibited predominantly filamentous morphology (∼70%) and fewer than 5% contained detectable post-F.

### Coverslip functionalization and virus immobilization

Coverslips for virus immobilization and imaging were prepared following a pegylation protocol modified from Piehler et al.^72^. Briefly, glass coverslips were cleaned with sonication using a 50% ethanol / 50% 3M NaOH solution for 30 minutes, followed by two rinses in 1L beakers of milliQ water. Coverslips were then cleaned using piranha solution (60% sulfuric acid, 40% H_2_O_2_) and sonication for 45 minutes, rinsed, dried, and functionalized with (3-glycidyloxypropyl)trimethoxysilane (GOPTS) for 1h at 75°C. Excess GOPTS was rinsed from coverslips using anhydrous acetone, and a mixture of biotin-PEG-amine / methoxy-PEG-amine (Rapp Polymere) was prepared in anhydrous acetone at a ratio of 10 mol% biotinylated PEG. The PEG solution was coupled to coverslips overnight at 75°C, rinsed twice in 1L beakers of milliQ water, and stored in milliQ water at 4°C until use.

For virus immobilization, pegylated coverslips were rinsed in ethanol, dried, and sealed with custom chambers made of polydimethylsiloxane with wells shaped using a 4mm biopsy punch. Wells were filled with PBS and incubated successively with streptavidin (5ug/ml in PBS) and anti-G antibody 3D3^27^ with a biotin site-specifically conjugated to the C-terminus of the heavy chain (see *Antibody cloning, expression, purification, and labeling*). After washing wells ten times with PBS to remove excess antibody, coverslips were stored at 4°C in a humidified enclosure for <2 days, until ready for use.

### Antibody cloning, expression, purification, and labelling

Antibody sequences used in this work are listed in Supporting information. VH and VL sequences were cloned into a human IgG1 backbone with a C-terminal ybbR tag using Gibson assembly. Verified clones were used to transfect T75 flasks of HEK293s at ∼85% confluency. At ∼12h post transfection, cells were washed twice with PBS to remove any residual IgG from the serum-containing culture media and grown for an addition 6 days in serum-free OptiMEM. Media containing secreted mAbs was collected and centrifuged at 1000xg to remove detached cells before purification with Protein G resin. Fab fragments (ADI-14359) and 5C4 IgM were expressed and purified analogously, with the exception that a C-terminal His(6)-tag on the heavy chains were used for affinity purification by Ni-NTA agarose in place of Protein G resin.

Eluted antibodies were quantified, diluted into a new buffer for enzymatic labeling (150mM NaCl, 25mM HEPES, 5mM MgCl_2_), and concentrated using centrifugal concentrators (VIVAspin 100K). Antibodies concentrated to ∼1mg/ml were then labeled overnight on ice using Sfp synthase and CoA-conjugated dyes, prepared as previously described^39^. Following removal of excess dye using PD-10 desalting columns, labeling efficiencies were determined spectrophotometrically to be >90% for all antibodies based on the number of heavy chains. For virus immobilization, we cloned and purified the anti-G antibody 3D3^27,73^ using the same protocol, but substituting CoA-biotin for the fluorescent dyes.

### C3/C4 deposition assay

C3 deposition assays were performed using IgG/IgM-depleted Normal Human Serum (NHS; Pelfreeze 34014). Fluorescent C3 and C4 were produced by labeling purified C3 or C4 (Complement Technologies, A113 and A105) using AF-488 dye functionalized with N-hydroxysuccinimide ester (Lumiprobe 11820). Labeling reactions were calibrated to prevent over-labeling of proteins and resulted in ∼0.6-1.0 dye molecules / protein, as determined via spectrophotometry. Complement reactions to monitor C3 deposition were prepared using complement buffer (150mM NaCl, 25mM HEPES, 0.5mM MgCl_2_, 0.15mM CaCl_2_) supplemented with 10mg/ml BSA, 5% IgG/IgM-depleted NHS, 50μg/ml 488-C3, and 10-20μg/ml fluorescent mAb (to assure rapid saturation of binding). Assuming a C3 concentration of 1mg/ml in NHS, approximately 30-50% of C3 in the experiment will carry a fluorophore. Prior to starting the reaction, all samples were washed thoroughly with complement buffer. Complement reactions were prepared on ice and added to virus samples before incubation at 37°C / 5% CO_2_ / 100% humidity for one hour. Following incubation, samples were washed three times with PBS to terminate the reaction and imaged immediately. A similar procedure was followed for timelapse experiments using C4, except NHS was used at a final concentration of 2.5% and 488-labeled C4 was supplemented at 10μg/ml. To synchronize recording with the start of the reaction, samples containing immobilized RSV were mounted in an incubated enclosure on the microscope prior to adding the complement components.

### C1 binding assay

C1 binding assays were performed using purified C1 (Complement Technologies A098) in the absence of other serum proteins and the presence of defined mAbs. C1 was diluted into complement buffer to a final concentration of 10μg/ml. The reaction also contained 10mg/ml BSA and saturating concentrations of defined mAbs (10-20μg/ml, as in C3 deposition assays). This mixture was incubated with enzymatically labeled RSV particles for 90 minutes at 4°C, washed three times in complement buffer, and labeled with an Alexa Fluor 488-conjugated anti-C1qA antibody (1A4; Santa Cruz sc-53544 AF488) at a 1:500 dilution in complement buffer for 30 minutes at room temperature before imaging.

### Fluorescence microscopy and image analysis

Following incubation with antibodies and complement components, we imaged opsonized RSV particles using a 60X 1.40 NA objective on a Nikon T2i microscope body equipped with a Yokogawa CSU-X spinning disk and ORCA-Flash4.0 V3 camera. For each sample condition per biological replicate, we collected images of ∼15 randomized fields of view, each containing ∼500-1000 RSV particles. The resulting image datasets were analyzed and plotted using custom Matlab scripts. Images were segmented using the F channel to identify pixels associated with each virus in the image. Following background subtraction, fluorescence intensities were integrated across all channels to obtain an integrated intensity for F, mAb, and C3/C4/C1 for each segmented particle. Integrated intensities were then plotted directly to determine population distributions (*e*.*g*. Figure 1C), or simplified further by determining the percentage of positive particles (*e*.*g*. Figure 1D).

### Virus photobleaching

Viruses labeled with Sulfo-Cy3 CLPMTGG via SrtA were immobilized on coverslips using antibodies against G (3D3) and imaged on an Olympus FluoView FV1200 laser scanning confocal microscope using a 60X 1.35 NA objective. A circular region ∼1μm in diameter that overlapped with a portion of the virus particle was selected and bleached using a 561nm laser, and fluorescence recovery was monitored by imaging at 5s intervals for one minute, including one image pre-bleach and one image immediately post-bleach. To identify globular particles enriched in post-F, a 488-labeled Fab fragment with VH and VL domains from ADI-14359 was added as a marker. Use of a Fab fragment for these experiments prevented antibody-mediated crosslinking of F that could alter fluorescence recovery. Time series of bleaching and recovery were used determine differences in F mobility. The percentage recovery was determined by generating an image mask from the difference between the first frame post-bleach and the last frame pre-bleach, to identify the bleached pixels. Intensities within the masked region were then integrated to quantify signal before bleaching, immediately after bleaching, and after a 20s recovery.

### Infectivity comparisons

Comparisons of virus infectivity following various treatments (Figure S1B, Figure S4A) used expression of the mTagBFP2 reporter to quantify infected cells. For comparisons of infectivity with or without fluorophores conjugated to F, viruses were labeled at 60hpi as described under *Collecting and labeling viruses*. Control samples were incubated at room temperature in parallel with labeled samples, and washed in the same way, to assure that collected viruses in all cases were shed exclusively over a two-hour period. 5μl of these samples were used to infect confluent A549 cells in 96-well plates, as described in *Preparing viruses*. Infection was quantified by counting BFP-positive cells across ∼10 fields of view at 10x magnification at 12hpi. Virus quantified in this way reflects only the particles shed from cells in a 2h period, and does not reflect the potentially substantial fraction of virus that remain cell-associated, or that were removed during wash steps.

For comparisons of untreated and osmotically-swollen viruses, 96-well plates containing infected A549 cells at 60hpi were washed with fresh media to remove older virus and returned to the incubator for 2h to allow new virus to shed. Collected samples were then split into experimental and control groups. For control groups, 5μl of shed virus was diluted into 95ul of 1x MEM with sodium bicarbonate and 7.5mM HEPES. For experimental groups, 5μl of shed virus was diluted into 75μl of 10mM HEPES (the low osmolarity buffer used for osmotic swelling), incubated for ∼1 minute, and added to 10μl of 10x sodium bicarbonate and 10x MEM, so that the final composition of control and experimental groups is matching. Samples were then used to infect confluent A549 cells in 96-well plates seeded the previous day. Infection was quantified by counting BFP-positive cells across ∼10 fields of view at 10x magnification at 12hpi.

### Creating polyclonal A549 CD55 and CD46 knock-out lines

A549 knockout cells were generated through transduction with lentivirus generated from the lentiCRISPR v2 packaging plasmid^74^. Three sgRNA sequences were selected using CRISPR KO and the design rules described by Doench *et al*.^75^. These were tested in small scale via transient transfection in HEK293s and the sgRNAs that yielded the highest efficiency (determined via immunofluorescence) were selected for lentivirus preparation and infection into A549s. The spacer sequences used for CD55 and CD46 sgRNAs are 5’-GCACCACCACAAATTGACAA-3’ (for CD55) and 5’-<u>G</u>TTTGTGATCGGAATCATACA-3’ (for CD46; underline indicates a nucleotide added for efficient transcription initiation). Polyclonal knockout cells were further enriched at the Washington University Flow Cytometry core, using a FACS Aria II to isolate cells negative for surface staining with fluorescent antibodies against CD46 (clone TRA-2-10) or CD55 (clone JS11).

### Modeling virus curvature

RSV particles are modeled in two simplified morphological states: filamentous particles - consisting of a cylindrical region of length *L* and radius *a*_*f*_ and two hemispherical caps - and globular particles, which we approximate as a sphere of radius *a*_*s*_. During the transition from a filament to a sphere, the surface area of the virus remains constant; this is constrained by the number of lipids packaged during assembly and the inability of lipid membranes to withstand area strains above ∼5%^76^. Conversely, the volume of the virus may change due to a flux of water into or out of the particle. The relationships in Figure S5A were obtained by applying a constant area constraint and equating the two surface areas (filament and sphere) and solving for the mean curvature in both cases.

## Acknowledgements

The authors would like to acknowledge members of the Vahey Lab for feedback and technical consultation and Dr. Ali Ellebedy for the antibody expression plasmid backbone. This work was supported by a Burroughs Wellcome Fund Career Awards at the Scientific Interfaces Grant and unrestricted funds from Washington University.

---

**Figure S1:**
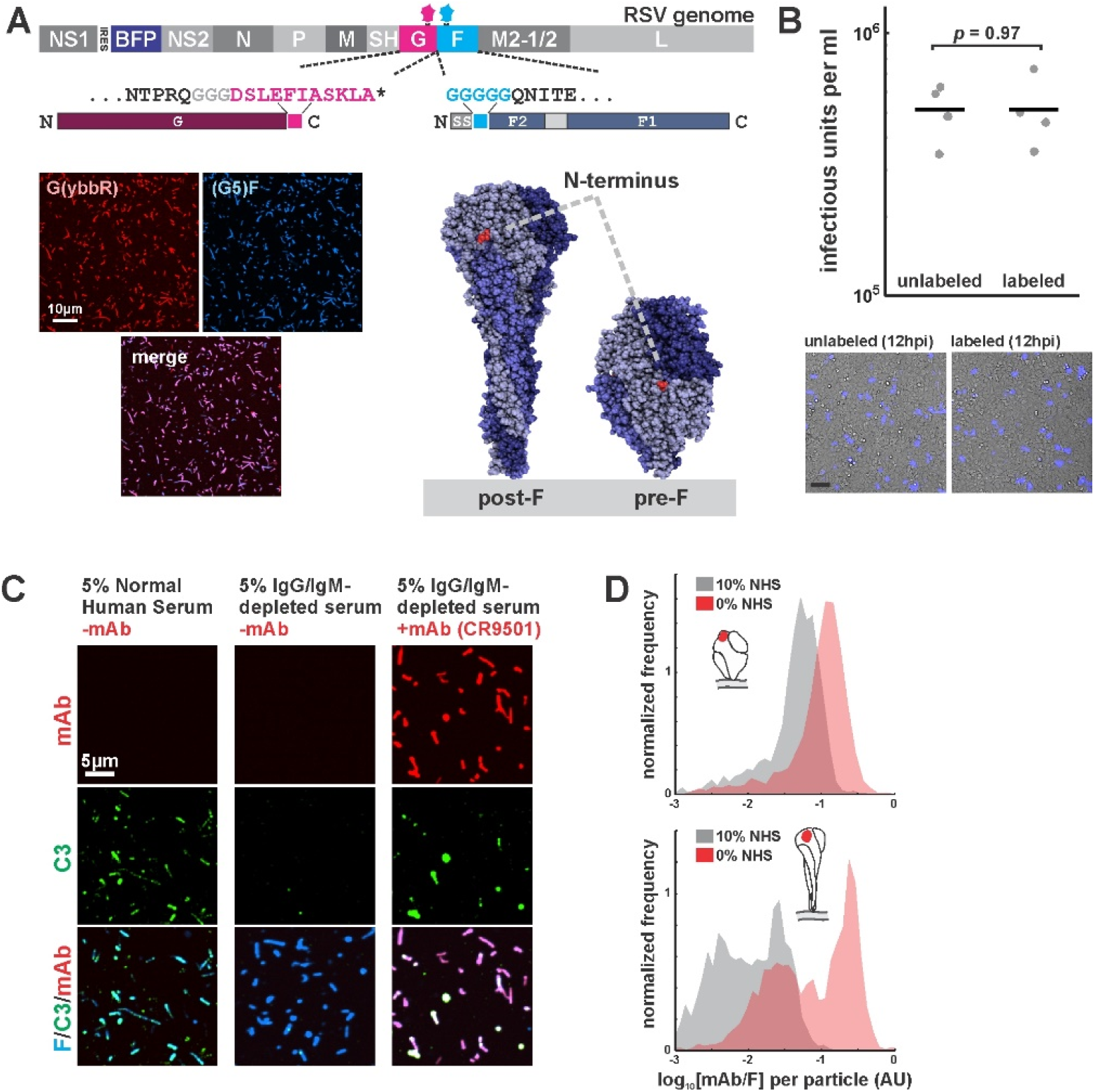
A fluorescence imaging-based approach to studying complement activation by RSV. (A) *Top*: Schematic of the RSV genome highlighting specific modifications: mTagBFP2 reporter expressed from an IRES following NS1, and tags for site-specific labeling on G (C-terminal ybbR tag) and F (N-terminal SrtA tag). *Bottom left*: images of RSV particles with labeled G and F. *Bottom right*: structures of pre- and postfusion F highlighting the location of the N-terminal tag, which remains accessible for labeling on both structures. (B) Comparison of RSV infectivity with and without fluorescent modifications to F. Infected cells were quantified at 12hpi by counting cells expressing the BFP reporter (representative images in lower panels; scale bar = 100μm). RSV particles quantified were those shed into the culture media during a 2h incubation and does not include cell-associated virus. Points show results from individual replicates; lines show mean values. *P*-value determined by a two-sample t-test. (C) Comparison of C3 deposition in complete normal human serum (5%) and IgG/IgM-depleted serum (5%), with and without supplemental mAbs. Images are displayed at matching contrast across each channel. (D) Antibody competition between fluorescent mAbs and polyclonal antibodies from normal human serum without IgG/IgM depletion. Results are shown as distributions of integrated mAb intensity normalized to integrated F intensity per RSV particle. *Top*: comparison of binding by pre-F-specific mAb 5C4. *Bottom*: comparison of binding by post-F-specific mAb ADI-14359 for viruses converted to post-F form by incubation with low ionic strength buffer (see also Figure 1B & E).

**Figure S2:**
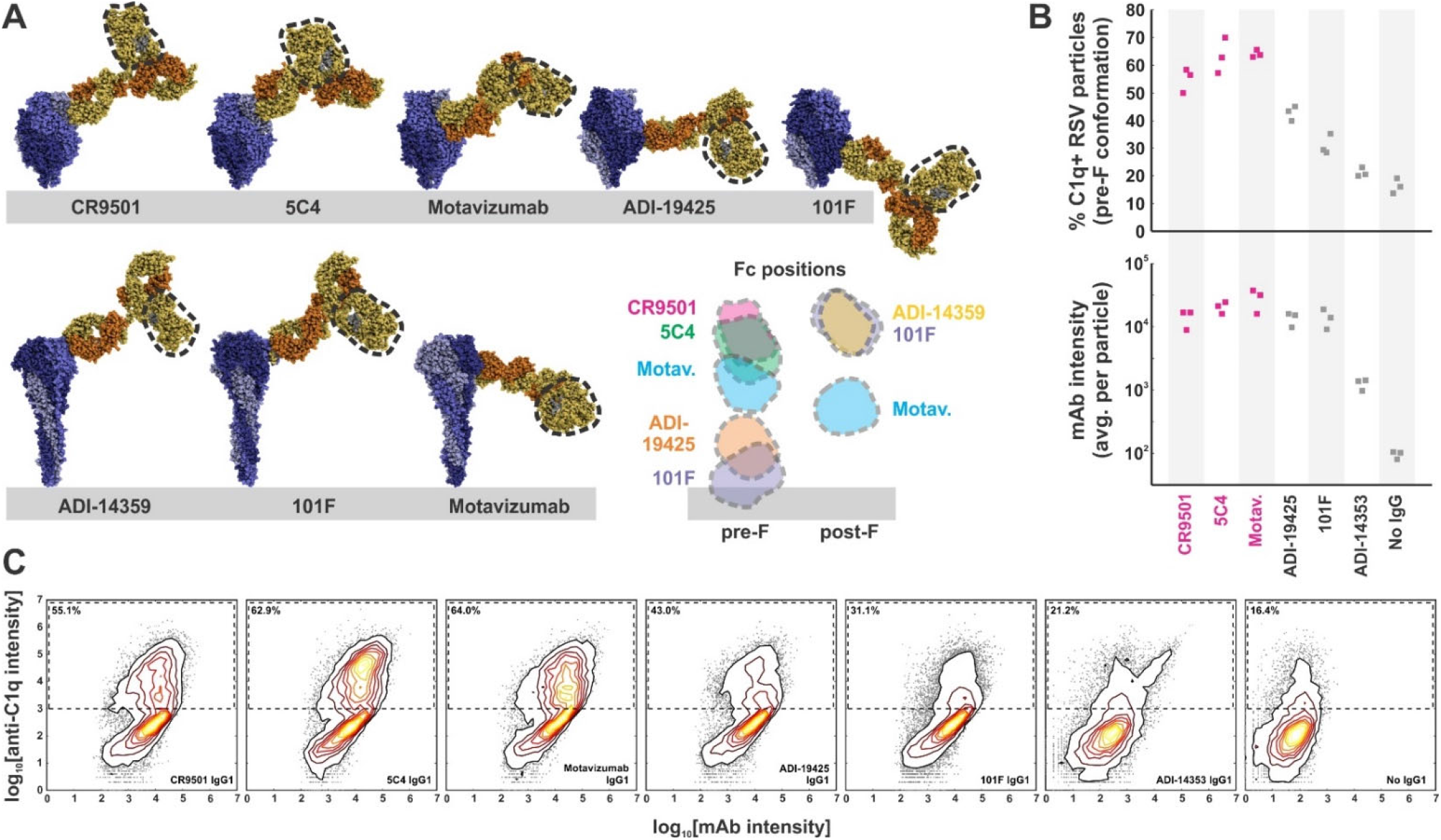
Complement activation and C1q binding varies with Fc position. (A) Modeling Fc positions for F-specific mAbs. Structures for RSV F (or portions thereof) with bound antibodies (PDB IDs 6OE4, 5W23, 4ZYP, 3IXT, 6APD, 6APB, and 3O45) were aligned with human IgG1 (PDB ID 1HZH) to determine representative locations accessible to antibody Fc regions (indicated by dashed outline). Distances from the viral membrane range from ∼1nm (101F) to ∼18nm (CR9501, ADI-14359). (B) C1q binding to predominantly pre-F RSV particles opsonized with different antibodies. The top plot shows the percentage of C1q+ particles, defined as those with a total intensity of anti-C1qA antibody >10^3^. The bottom plot shows the intensity of anti-F mAb for each condition. Individual points represent values for three biological replicates. Antibodies determined to activate complement from pre-F antigens (Figure 1) are shown in magenta. (C) Distributions of anti-F mAb intensities (horizontal axis) and anti-C1qA antibody intensities (vertical axis) for different anti-F mAbs bound to pre-F particles. Particles within the dashed rectangles indicate those that are C1q-positive, and the percentage of these particles is indicated in the upper left. Distributions are combined from the same three biological replicates represented in *B*.

**Figure S3:**
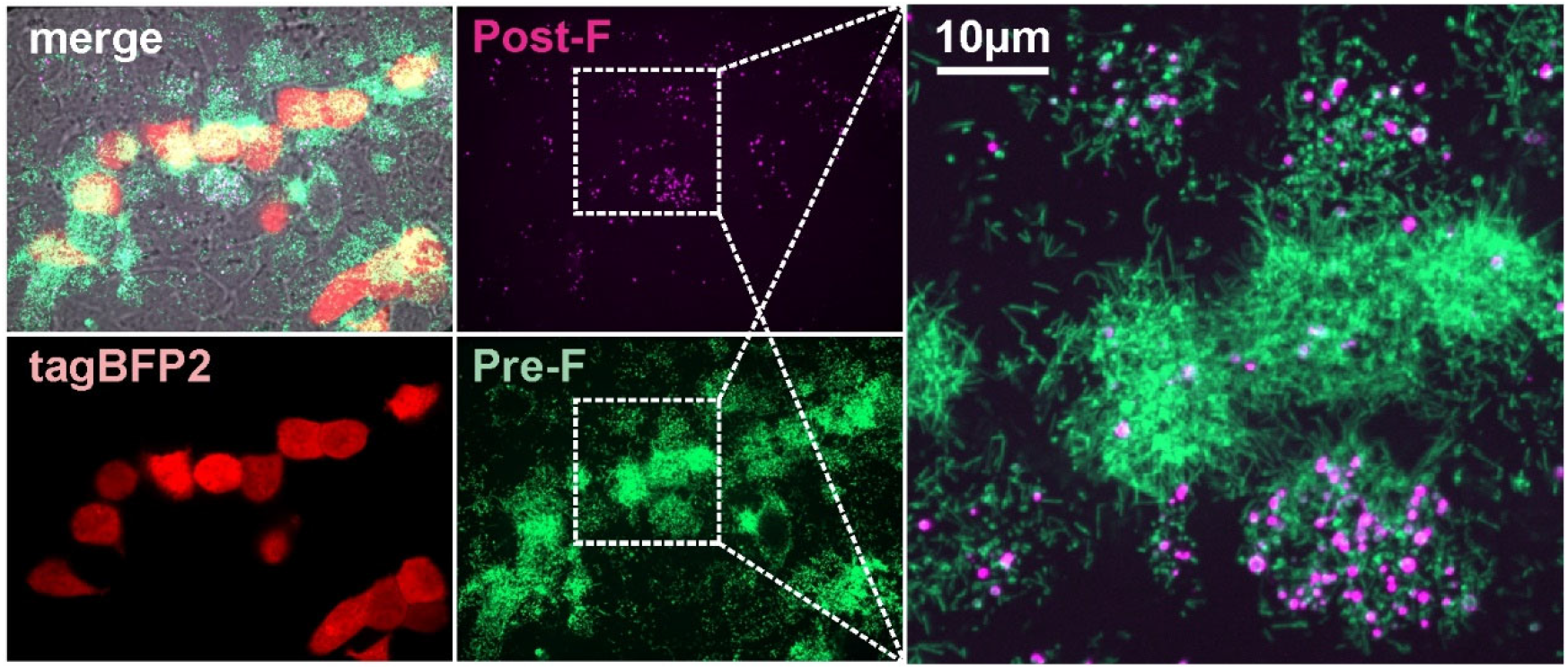
Pre-F and post-F containing RSV particles occur naturally in cell culture. Images of live RSV-infected A549 cells at 48hpi showing mTagBFP2 reporter (indicating infected cells) along with pre- and postfusion F, labeled using 5C4 (Alexa Fluor 488) and ADI-14359 (Alexa Fluor 555) mAbs added directly to culture medium. Images are displayed as maximum intensity projections from a three-dimensional confocal stack. Post-F enriched particles with globular morphology are present on the surface of both RSV-infected and neighboring uninfected cells.

**Figure S4:**
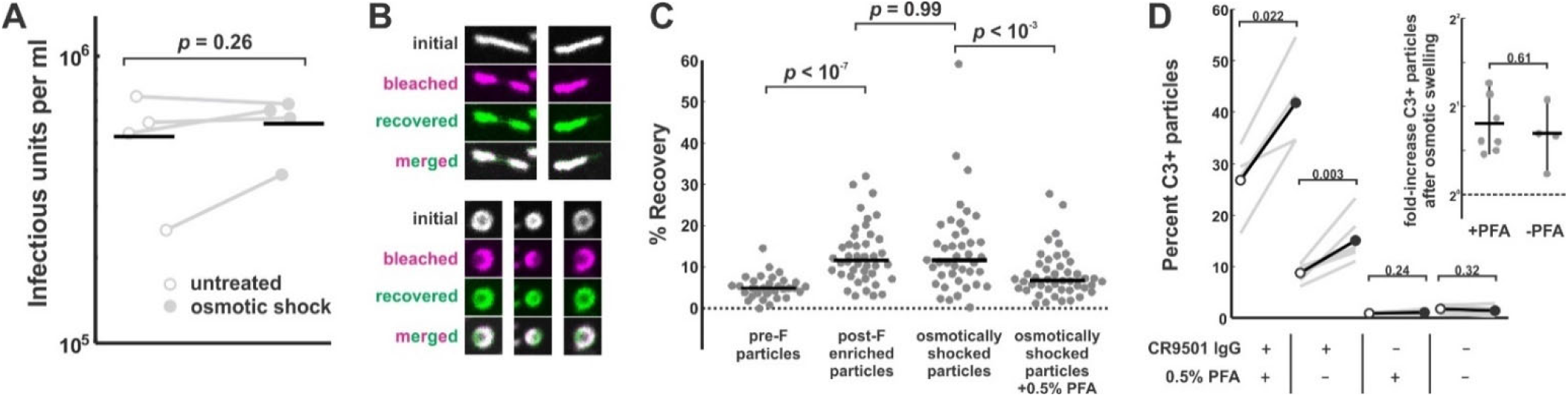
Osmotic swelling detaches the RSV matrix from the viral membrane with no loss of infectivity. (A) Comparison of infectivity (quantified as single-round infectious units per ml) of virus shed from A549 cells during a two-hour period starting at 60hpi. RSV samples were divided into control (untreated) and experimental groups (osmotic shock), and used to infect confluent monolayers of A549 cells. Infected cells were quantified at 12hpi by counting cells expressing the BFP reporter. The experiment does not account for RSV particles that remain cell-associated. (B) Images of photobleached / recovered RSV filaments (top) and spheres (bottom), where spheres are obtained through osmotic swelling. Magenta images show virus particles immediately after bleaching; green images show viruses after 20 seconds of recovery. (C) Quantification of fluorescence recovery of different RSV particle subsets. Points represent individual viruses and black lines represent median values. *P*-values are determined using a two-sample KS test. (D) C3 deposition on RSV particles with (filled circles) and without (open circles) osmotic swelling. Gray lines connecting points indicate paired biological replicates. Fixation with 0.5% PFA was used to restrict antigen mobility, as shown in *C. Inset*: fold-change in percent C3-positive particles following osmotic swelling, with and without treatment with 0.5% PFA.

**Figure S5:**
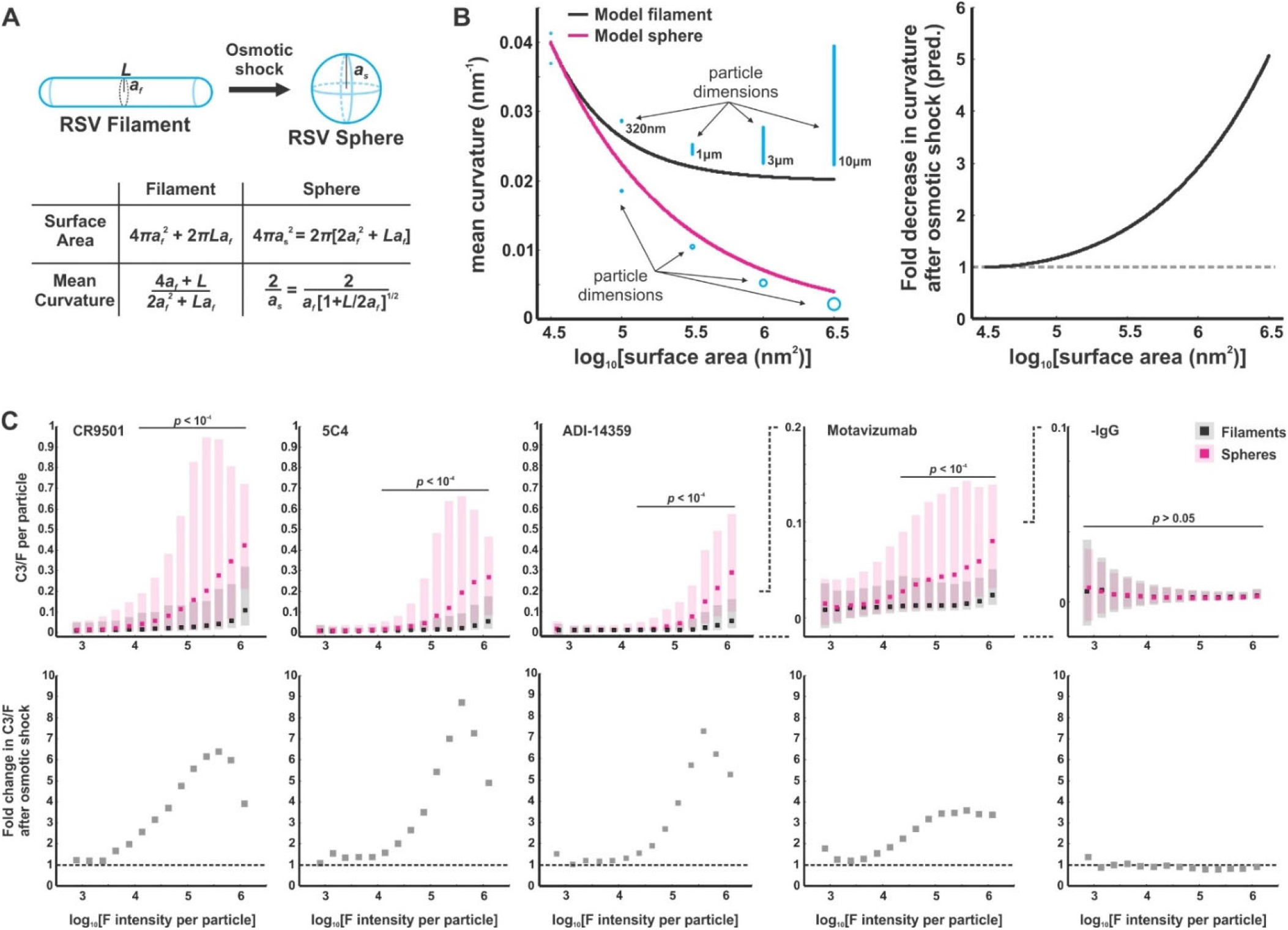
C3 deposition on RSV particles increases with decreasing particle curvature. (A) Model of an idealized morphological transformation in RSV. Detachment of the viral matrix leads to the rounding of virus particles. Mean curvature can be predicted from particle geometry and the constraint that surface area is conserved during the shape transformation. (B) *Left*: plot of the relationships in *A* showing mean curvature versus particle surface area. Schematics of viral filaments / spheres are drawn approximately to scale in blue. *Right*: relationships from the left plot, displayed as the ratio of filament mean curvature to sphere mean curvature as a function of surface area. (C) *Top*: data for C3 deposition driven by four activating antibodies (CR9501, 5C4, and Motavizumab for pre-F and ADI-14359 for post-F) and a negative control with no antibody. Black points/bars show median / 25^th^-75^th^ percentile data for viruses preserved in the filamentous state and magenta points/bars show data for osmotically swollen viruses. *Bottom*: same data as in the top plots, displayed as fold-increase in median C3/F for spherical vs. filamentous morphological states.

## Notes

### Competing Interest Statement

The authors have declared no competing interest.

## References

1. O’Brien, K. B., Morrison, T. E., Dundore, D. Y., Heise, M. T. & Schultz-Cherry, S. A protective role for complement C3 protein during pandemic 2009 H1N1 and H5N1 influenza A virus infection. PLoS One 6, e17377 (2011).

2. Kopf, M., Abel, B., Gallimore, A., Carroll, M. & Bachmann, M. F. Complement component C3 promotes T-cell priming and lung migration to control acute influenza virus infection. Nat Med 8, 373–378 (2002).

3. Bottermann, M. et al.. Complement C4 Prevents Viral Infection through Capsid Inactivation. Cell Host & Microbe 25, 617-629.e7 (2019).

4. Bladen, H. A., Evans, R. T. & Mergenhagen, S. E. Lesions in Escherichia coli membranes after action of antibody and complement. Journal of Bacteriology 91, 2377–2381 (1966).

5. Mishra, M. et al.. Pseudomonas aeruginosa Psl polysaccharide reduces neutrophil phagocytosis and the oxidative response by limiting complement-mediated opsonization. Cell Microbiol 14, 95–106 (2012).

6. Roestenberg, M. et al.. Complement activation in experimental human malaria infection. Transactions of the Royal Society of Tropical Medicine and Hygiene 101, 643–649 (2007).

7. Engstler, M. et al.. Hydrodynamic Flow-Mediated Protein Sorting on the Cell Surface of Trypanosomes. Cell 131, 505–515 (2007).

8. Kurtovic, L. et al.. Human antibodies activate complement against Plasmodium falciparum sporozoites, and are associated with protection against malaria in children. BMC Med 16, 61 (2018).

9. Dunkelberger, J. R. & Song, W.-C. Complement and its role in innate and adaptive immune responses. Cell Res 20, 34–50 (2010).

10. Ochsenbein, A. F. et al.. Protective T Cell–Independent Antiviral Antibody Responses Are Dependent on Complement. Journal of Experimental Medicine 190, 1165–1174 (1999).

11. Phan, T. G., Grigorova, I., Okada, T. & Cyster, J. G. Subcapsular encounter and complement-dependent transport of immune complexes by lymph node B cells. Nat Immunol 8, 992–1000 (2007).

12. Reynes, M. et al.. Human follicular dendritic cells express CR1, CR2, and CR3 complement receptor antigens. J Immunol 135, 2687–2694 (1985).

13. Hebell, T., Ahearn, J. & Fearon, D. Suppression of the immune response by a soluble complement receptor of B lymphocytes. Science 254, 102–105 (1991).

14. Dempsey, P. W., Allison, M. E. D., Akkaraju, S., Goodnow, C. C. & Fearon, D. T. C3d of Complement as a Molecular Adjuvant: Bridging Innate and Acquired Immunity. Science 271, 348– 350 (1996).

15. Lukácsi, S., Nagy-Baló, Z., Erdei, A., Sándor, N. & Bajtay, Z. The role of CR3 (CD11b/CD18) and CR4 (CD11c/CD18) in complement-mediated phagocytosis and podosome formation by human phagocytes. Immunology Letters 189, 64–72 (2017).

16. Cornacoff, J. B. et al.. Primate erythrocyte-immune complex-clearing mechanism. J. Clin. Invest. 71, 236–247 (1983).

17. Bera, M. M. et al.. Th17 cytokines are critical for respiratory syncytial virus-associated airway hyperreponsiveness through regulation by complement C3a and tachykinins. J Immunol 187, 4245– 4255 (2011).

18. Peng, Q., Li, K., Sacks, S. & Zhou, W. The Role of Anaphylatoxins C3a and C5a in Regulating Innate and Adaptive Immune Responses. IADT 8, 236–246 (2009).

19. Gralinski, L. E. et al.. Complement Activation Contributes to Severe Acute Respiratory Syndrome Coronavirus Pathogenesis. mBio 9, e01753-18, /mbio/9/5/mBio.01753-18.atom (2018).

20. Ricklin, D., Reis, E. S. & Lambris, J. D. Complement in disease: a defence system turning offensive. Nat Rev Nephrol 12, 383–401 (2016).

21. Corbeil, S., Seguin, C. & Trudel, M. Involvement of the complement system in the protection of mice from challenge with respiratory syncytial virus Long strain following passive immunization with monoclonal antibody 18A2B2. Vaccine 14, 521–525 (1996).

22. Bukreyev, A., Yang, L. & Collins, P. L. The Secreted G Protein of Human Respiratory Syncytial Virus Antagonizes Antibody-Mediated Restriction of Replication Involving Macrophages and Complement. Journal of Virology 86, 10880–10884 (2012).

23. Polack, F. P. et al.. A Role for Immune Complexes in Enhanced Respiratory Syncytial Virus Disease. Journal of Experimental Medicine 196, 859–865 (2002).

24. McLellan, J. S. et al.. Structure of RSV fusion glycoprotein trimer bound to a prefusion-specific neutralizing antibody. Science 340, 1113–1117 (2013).

25. Fedechkin, S. O., George, N. L., Wolff, J. T., Kauvar, L. M. & DuBois, R. M. Structures of respiratory syncytial virus G antigen bound to broadly neutralizing antibodies. Sci. Immunol. 3, eaar3534 (2018).

26. Gilman, M. S. A. et al.. Rapid profiling of RSV antibody repertoires from the memory B cells of naturally infected adult donors. Science Immunology 1, eaaj1879–eaaj1879 (2016).

27. Collarini, E. J. et al.. Potent High-Affinity Antibodies for Treatment and Prophylaxis of Respiratory Syncytial Virus Derived from B Cells of Infected Patients. J Immunol 183, 6338–6345 (2009).

28. Battles, M. B. & McLellan, J. S. Respiratory syncytial virus entry and how to block it. Nat Rev Microbiol 17, 233–245 (2019).

29. Griffiths, C. D. et al.. IGF1R is an entry receptor for respiratory syncytial virus. Nature 583, 615–619 (2020).

30. Johnson, S. M. et al.. Respiratory Syncytial Virus Uses CX3CR1 as a Receptor on Primary Human Airway Epithelial Cultures. PLoS Pathog 11, e1005318 (2015).

31. Chirkova, T. et al.. CX3CR1 is an important surface molecule for respiratory syncytial virus infection in human airway epithelial cells. J Gen Virol 96, 2543–2556 (2015).

32. Zhivaki, D. et al.. Respiratory Syncytial Virus Infects Regulatory B Cells in Human Neonates via Chemokine Receptor CX3CR1 and Promotes Lung Disease Severity. Immunity 46, 301–314 (2017).

33. Graham, B. S. Vaccine development for respiratory syncytial virus. Current Opinion in Virology 23, 107–112 (2017).

34. Goodwin, E. et al.. Infants Infected with Respiratory Syncytial Virus Generate Potent Neutralizing Antibodies that Lack Somatic Hypermutation. Immunity 48, 339-349.e5 (2018).

35. McLellan, J. S. et al.. Structure-Based Design of a Fusion Glycoprotein Vaccine for Respiratory Syncytial Virus. Science 342, 592–598 (2013).

36. Marcandalli, J. et al.. Induction of Potent Neutralizing Antibody Responses by a Designed Protein Nanoparticle Vaccine for Respiratory Syncytial Virus. Cell 176, 1420-1431.e17 (2019).

37. Crank, M. C. et al.. A proof of concept for structure-based vaccine design targeting RSV in humans. Science 365, 505–509 (2019).

38. Acosta, P. L., Caballero, M. T. & Polack, F. P. Brief History and Characterization of Enhanced Respiratory Syncytial Virus Disease. Clinical and Vaccine Immunology 23, 189–195 (2016).

39. Vahey, M. D. & Fletcher, D. A. Low-Fidelity Assembly of Influenza A Virus Promotes Escape from Host Cells. Cell 176, 281-294.e19 (2019).

40. Yin, J., Lin, A. J., Golan, D. E. & Walsh, C. T. Site-specific protein labeling by Sfp phosphopantetheinyl transferase. Nature Protocols 1, 280–285 (2006).

41. Theile, C. S. et al.. Site-specific N-terminal labeling of proteins using sortase-mediated reactions. Nature Protocols 8, 1800–1807 (2013).

42. Gilman, M. S. A. et al.. Transient opening of trimeric prefusion RSV F proteins. Nat Commun 10, 2105 (2019).

43. Wu, H. et al.. Development of motavizumab, an ultra-potent antibody for the prevention of respiratory syncytial virus infection in the upper and lower respiratory tract. J Mol Biol 368, 652–665 (2007).

44. Wu, S.-J. et al.. Characterization of the epitope for anti-human respiratory syncytial virus F protein monoclonal antibody 101F using synthetic peptides and genetic approaches. J Gen Virol 88, 2719–2723 (2007).

45. Hicks, S. N. set al. Five Residues in the Apical Loop of the Respiratory Syncytial Virus Fusion Protein F _2_ Subunit Are Critical for Its Fusion Activity. J Virol 92, e00621–18, /jvi/92/15/e00621-18.atom (2018).

46. Chaiwatpongsakorn, S., Epand, R. F., Collins, P. L., Epand, R. M. & Peeples, M. E. Soluble Respiratory Syncytial Virus Fusion Protein in the Fully Cleaved, Pretriggered State Is Triggered by Exposure to Low-Molarity Buffer. Journal of Virology 85, 3968–3977 (2011).

47. Diebolder, C. A. et al. Complement Is Activated by IgG Hexamers Assembled at the Cell Surface. Science 343, 1260–1263 (2014).

48. Ugurlar, D. et al. Structures of C1-IgG1 provide insights into how danger pattern recognition activates complement. Science 359, 794–797 (2018).

49. Wang, G. et al. Molecular Basis of Assembly and Activation of Complement Component C1 in Complex with Immunoglobulin G1 and Antigen. Mol Cell 63, 135–145 (2016).

50. Sharp, T. H. et al. Insights into IgM-mediated complement activation based on in situ structures of IgM-C1-C4b. Proc Natl Acad Sci USA 201901841 (2019) doi:10.1073/pnas.1901841116.

51. Li, Y. & Parks, G. Relative Contribution of Cellular Complement Inhibitors CD59, CD46, and CD55 to Parainfluenza Virus 5 Inhibition of Complement-Mediated Neutralization. Viruses 10, 219 (2018).

52. Hutchinson, E. C. et al. Conserved and host-specific features of influenza virion architecture. Nature Communications 5, 4816 (2014).

53. Marschang, P., Sodroski, J., Würzner, R. & Dierich, M. P. Decay-accelerating factor (CD55) protects human immunodeficiency virus type 1 from inactivation by human complement. Eur. J. Immunol. 25, 285–290 (1995).

54. Noris, M. & Remuzzi, G. Overview of Complement Activation and Regulation. Seminars in Nephrology 33, 479–492 (2013).

55. Liljeroos, L., Krzyzaniak, M. A., Helenius, A. & Butcher, S. J. Architecture of respiratory syncytial virus revealed by electron cryotomography. Proceedings of the National Academy of Sciences 110, 11133–11138 (2013).

56. Ke, Z. et al. The Morphology and Assembly of Respiratory Syncytial Virus Revealed by Cryo-Electron Tomography. Viruses 10, 446 (2018).

57. Förster, A., Maertens, G. N., Farrell, P. J. & Bajorek, M. Dimerization of Matrix Protein Is Required for Budding of Respiratory Syncytial Virus. J. Virol. 89, 4624–4635 (2015).

58. Shaikh, F. Y. et al. A Critical Phenylalanine Residue in the Respiratory Syncytial Virus Fusion Protein Cytoplasmic Tail Mediates Assembly of Internal Viral Proteins into Viral Filaments and Particles. mBio 3, e00270–11 (2012).

59. Pedersen, M. B. et al. Curvature of Synthetic and Natural Surfaces Is an Important Target Feature in Classical Pathway Complement Activation. J.I. 184, 1931–1945 (2010).

60. Zeuthen, C. M. et al. C1q recognizes antigen-bound IgG in a curvature-dependent manner. Nano Res. 13, 1651–1658 (2020).

61. Westas Janco, E., Hulander, M. & Andersson, M. Curvature-dependent effects of nanotopography on classical immune complement activation. Acta Biomaterialia 74, 112–120 (2018).

62. Wasilewski, S., Calder, L. J., Grant, T. & Rosenthal, P. B. Distribution of surface glycoproteins on influenza A virus determined by electron cryotomography. Vaccine 30, 7368–7373 (2012).

63. Peukes, J. et al. The native structure of the assembled matrix protein 1 of influenza A virus. Nature 587, 495–498 (2020).

64. Ke, Z. et al. Promotion of virus assembly and organization by the measles virus matrix protein. Nat Commun 9, 1736 (2018).

65. Beniac, D. R. & Booth, T. F. Structure of the Ebola virus glycoprotein spike within the virion envelope at 11?Å resolution. Sci Rep 7, 46374 (2017).

66. Doorduijn, D. J. et al. Bacterial killing by complement requires direct anchoring of membrane attack complex precursor C5b-7. PLoS Pathog 16, e1008606 (2020).

67. Liljeroos, L., Huiskonen, J. T., Ora, A., Susi, P. & Butcher, S. J. Electron cryotomography of measles virus reveals how matrix protein coats the ribonucleocapsid within intact virions. Proceedings of the National Academy of Sciences 108, 18085–18090 (2011).

68. Battisti, A. J. et al. Structure and assembly of a paramyxovirus matrix protein. Proceedings of the National Academy of Sciences 109, 13996–14000 (2012).

69. Hotard, A. L. et al. A stabilized respiratory syncytial virus reverse genetics system amenable to recombination-mediated mutagenesis. Virology 434, 129–136 (2012).

70. Warming, S. Simple and highly efficient BAC recombineering using galK selection. Nucleic Acids Research 33, e36–e36 (2005).

71. Vahey, M. D. & Fletcher, D. A. Influenza A virus surface proteins are organized to help penetrate host mucus. eLife 8, (2019).

72. Piehler, J., Brecht, A., Valiokas, R., Liedberg, B. & Gauglitz, G. A high-density poly(ethylene glycol) polymer brush for immobilization on glass-type surfaces. Biosensors and Bioelectronics 15, 473–481 (2000).

73. Fedechkin, S. O., George, N. L., Wolff, J. T., Kauvar, L. M. & DuBois, R. M. Structures of respiratory syncytial virus G antigen bound to broadly neutralizing antibodies. Sci. Immunol. 3, eaar3534 (2018).

74. Sanjana, N. E., Shalem, O. & Zhang, F. Improved vectors and genome-wide libraries for CRISPR screening. Nat Methods 11, 783–784 (2014).

75. Doench, J. G. et al. Optimized sgRNA design to maximize activity and minimize off-target effects of CRISPR-Cas9. Nat Biotechnol 34, 184–191 (2016).

76. Needham, D. & Nunn, R. S. Elastic deformation and failure of lipid bilayer membranes containing cholesterol. Biophysical Journal 58, 997–1009 (1990).

