## Supporting Information for "A morphological transformation in respiratory syncytial virus leads to enhanced complement activation"

### RSV sequences

**RSV A2 (Line 19F):**

NS1-IRES-mTagBFP2/NS2/NP/P/M/SH/G(**ybbR**)/(**G5**)F/M2/L

ACGCGAAAAAATGCGTACAACAAACTTGCATAAACCAAAAAAATGGGGCAAATAAGAATTTGATAAGTACCACTTAAATTTAACTCCCTTGGTTAGAGATGGGCAGCAATTC
ATTGAGTATGATAAAAGTTAGATTACAAAATTTGTTTGACAATGATGAAGTAGCATTGTTAAAAATAACATGCTATACTGATAAATTAATACATTTAACTAACGCTTTGGCT
AAGGCAGTGATACATACAATCAAATTGAATGGCATTGTGTTTGTGCATGTTATTACAAGTAGTGATATTTGCCCTAATAATAATATTGTAGTAAAATCCAATTTCACAACAA
TGCCAGTACTACAAAATGGAGGTTATATATGGGAAATGATGGAATTAACACATTGCTCTCAACCTAATGGTCTACTAGATGACAATTGTGAAATTAAATTCTCCAAAAAACT
AAGTGATTCAACAATGACCAATTATATGAATCAATTATCTGAATTACTTGGATTTGATCTTAATCCATAACGGTTTCCCTCTAGCGGGATCAATTCCGCCCCCCCCCCCTAA
CGTTACTGGCCGAAGCCGCTTGGAATAAGGCCGGTGTGCGTTTGTCTATATGTTATTTTCCACCATATTGCCGTCTTTTGGCAATGTGAGGGCCCGGAAACCTGGCCCTGTC
TTCTTGACGAGCATTCCTAGGGGTCTTTCCCCTCTCGCCAAAGGAATGCAAGGTCTGTTGAATGTCGTGAAGGAAGCAGTTCCTCTGGAAGCTTCTTGAAGACAAACAACGT
CTGTAGCGACCCTTTGCAGGCAGCGGAACCCCCCACCTGGCGACAGGTGCCTCTGCGGCCAAAAGCCACGTGTATAAGATACACCTGCAAAGGCGGCACAACCCCAGTGCCA
CGTTGTGAGTTGGATAGTTGTGGAAAGAGTCAAATGGCTCTCCTCAAGCGTATTCAACAAGGGGCTGAAGGATGCCCAGAAGGTACCCCATTGTATGGGATCTGATCTGGGG
CCTCGGTGCACATGCTTTACATGTGTTTAGTCGAGGTTAAAAAAACGTCTAGGCCCCCCGAACCACGGGGACGTGGTTTTCCTTTGAAAAACACGATAATACCATGAGCGAG
CTGATTAAGGAGAACATGCACATGAAGCTGTACATGGAGGGCACCGTGGACAACCATCACTTCAAGTGCACATCCGAGGGCGAAGGCAAGCCCTACGAGGGCACCCAGACCA
TGAGAATCAAGGTGGTCGAGGGCGGCCCTCTCCCCTTCGCCTTCGACATCCTGGCTACTAGCTTCCTCTACGGCAGCAAGACCTTCATCAACCACACCCAGGGCATCCCCGA
CTTCTTCAAGCAGTCCTTCCCTGAGGGCTTCACATGGGAGAGAGTCACCACATACGAAGACGGGGGCGTGCTGACCGCTACCCAGGACACCAGCCTCCAGGACGGCTGCCTC
ATCTACAACGTCAAGATCAGAGGGGTGAACTTCACATCCAACGGCCCTGTGATGCAGAAGAAAACACTCGGCTGGGAGGCCTTCACCGAGACGCTGTACCCCGCTGACGGCG
GCCTGGAAGGCAGAAACGACATGGCCCTGAAGCTCGTGGGCGGGAGCCATCTGATCGCAAACGCCAAGACCACATATAGATCCAAGAAACCCGCTAAGAACCTCAAGATGCC
TGGCGTCTACTATGTGGACTACAGACTGGAAAGAATCAAGGAGGCCAACAACGAGACCTACGTCGAGCAGCACGAGGTGGCAGTGGCCAGATACTGCGACCTCCCTAGCAAA
CTGGGGCACAAGCTTAATTAAATTATAATTAATATCAACTAGCAAATCAATGTCACTAACACCATTAGTTAATATAAAACTTAACAGAAGACAAAAATGGGGCAAATAAATC
AATTCAGCCAACCCAACCATGGACACAACCCACAATGATAATACCACCACAAAGACTGATGATCACAGACATGAGACCGTTGTCACTTGAGACCATAATAACATCACTAACCA
GAGACATCATAACACACAAATTTATATACTTGATAAATCATGAATGCATAGTGAGAAAACTTGATGAAAAACAGGCCACATTTACATTCCTGGTCAACTATGAAATGAAACT
ATTACACAAAGTAGGAAGCACTAAATATAAAAAATATACTGAATACAACACAAAATATGGCACTTTCCCTATGCCAATATTCATCAATCATGATGGGTTCTTAGAATGCATT
GGCATTAAGCCTACAAAGCATACTCCCATAATATACAAGTATGATCTCAATCCATAAATTTCAACACAATATTCACACAATCTAAAACAACAACTCTATGCATAACTATACT
CCATAGTCCAGATGGAGCCTGAAAATTATAGTAATTTAAAATTAAGGAGAGATATAAGATAGAAGATGGGGCAAATACAAAGATGGCTCTTAGCAAAGTCAAGTTGAATGAT
ACACTCAACAAAGATCAACTTCTGTCATCCAGCAAATACACCATCCAACGGAGCACAGGAGATAGTATTGATACTCCTAATTATGATGTGCAGAAACACATCAATAAGTTAT
GTGGCATGTTATTAATCACAGAAGATGCTAATCATAAATTCACTGGGTTAATAGGTATGTTATATGCGATGTCTAGGTTAGGAAGAGAAGACACCATAAAAATACTCAGAGA
TGCGGGATATCATGTAAAAGCAAATGGAGTAGATGTAACAACACATCGTCAAGACATTAATGGAAAAGAAATGAAATTTGAAGTGTTAACATTGGCAAGCTTAACAACTGAA
ATTCAAATCAACATTGAGATAGAATCTAGAAAATCCTACAAAAAAATGCTAAAAGAAATGGGAGAGGTAGCTCCAGAATACAGGCATGACTCTCCTGATTGTGGGATGATAA
TATTATGTATAGCAGCATTAGTAATAACTAAATTAGCAGCAGGGGACAGATCTGGTCTTACAGCCGTGATTAGGAGAGCTAATAATGTCCTAAAAAATGAAATGAAACGTTA
CAAAGGCTTACTACCCAAGGACATAGCCAACAGCTTCTATGAAGTGTTTGAAAAACATCCCCACTTTATAGATGTTTTTGTTCATTTTGGTATAGCACAATCTTCTACCAGA
GGTGGCAGTAGAGTTGAAGGGATTTTTGCAGGATTGTTTATGAATGCCTATGGTGCAGGGCAAGTGATGTTACGGTGGGGAGTCTTAGCAAAATCAGTTAAAAATATTATGT
TAGGACATGCTAGTGTGCAAGCAGAAATGGAACAAGTTGTTGAGGTTTATGAATATGCCCAAAAATTGGGTGGTGAAGCAGGATTCTACCATATATTGAACAACCCAAAAGC
ATCATTATTATCTTTGACTCAATTTCCTCACTTCTCCAGTGTAGTATTAGGCAATGCTGCTGGCCTAGGCATAATGGGAGAGTACAGAGGTACACCGAGGAATCAAGATCTA
TATGATGCAGCAAAGGCATATGCTGAACAACTCAAAGAAAATGGTGTGATTAACTACAGTGTACTAGACTTGACAGCAGAAGAACTAGAGGCTATCAAACATCAGCTTAATC
CAAAAGATAATGATGTAGAGCTTTGAGTTAATAAAAAATGGGGCAAATAAATCATCATGGAAAAGTTTGCTCCTGAATTCCATGGAGAAGATGCAAACAACAGGGCTACTAA
ATTCCTAGAATCAATAAAGGGCAAATTCACATCACCCAAAGATCCCAAGAAAAAAGATAGTATCATATCTGTCAACTCAATAGATATAGAAGTAACCAAAGAAAGCCCTATA
ACATCAAATTCAACTATTATCAACCCAACAAATGAGACAGATGATACTGCAGGGAACAAGCCCAATTATCAAAGAAAACCTCTAGTAAGTTTCAAAGAAGACCCTACACCAA
GTGATAATCCCTTTTCTAAACTATACAAAGAAACCATAGAAACATTTGATAACAATGAAGAAGAATCCAGCTATTCATACGAAGAAATAAATGATCAGACAAACGATAATAT
AACAGCAAGATTAGATAGGATTGATGAAAAATTAAGTGAAATACTAGGAATGCTTCACACATTAGTAGTGGCAAGTGCAGGACCTACATCTGCTCGGGATGGTATAAGAGAT
GCCATGATTGGTTTAAGAGAAGAAATGATAGAAAAAATCAGAACTGAAGCATTAATGACCAATGACAGATTAGAAGCTATGGCAAGACTCAGGAATGAGGAAAGTGAAAAGA
TGGCAAAAGACACATCAGATGAAGTGTCTCTCAATCCAACATCAGAGAAATTGAACAACCTATTGGAAGGGAATGATAGTGACAATGATCTATCACTTGAAGATTTCTGATT
AGTTACCACTCTTCACATCAACACACAATACCAACAGAAGACCAACAAACTAACCAACCCAATCATCCAACCAAACATCCATCCGCCAATCAGCCAAACAGCCAACAAAACA
ACCAGCCAATCCAAAACTAACCACCCGGAAAAAATCTATAATATAGTTACAAAAAAAGGAAAGGGTGGGGCAAATATGGAAACATACGTGAACAAGCTTCACGAAGGCTCCA
CATACACAGCTGCTGTTCAATACAATGTCTTAGAAAAAGACGATGACCCTGCATCACTTACAATATGGGTGCCCATGTTCCAATCATCTATGCCAGCAGATTTACTTATAAA
AGAACTAGCTAATGTCAACATACTAGTGAAACAAATATCCACACCCAAGGGACCTTCACTAAGAGTCATGATAAACTCAAGAAGTGCAGTGCTAGCACAAATGCCCAGCAAA
TTTACCATATGCGCTAATGTGTCCTTGGATGAAAGAAGCAAACTAGCATATGATGTAACCACACCCTGTGAAATCAAGGCATGTAGTCTAACATGCCTAAAATCAAAAAATA
TGTTGACTACAGTTAAAGATCTCACTATGAAGACACTCAACCCTACACATGATATTATTGCTTTATGTGAATTTGAAAACATAGTAACATCAAAAAAAGTCATAATACCAAC
ATACCTAAGATCCATCAGTGTCAGAAATAAAGATCTGAACACACTTGAAAATATAACAACCACTGAATTCAAAAATGCTATCACAAATGCAAAAATCATCCCTTACTCAGGA
TTACTATTAGTCATCACAGTGACTGACAACAAAGGAGCATTCAAATACATAAAGCCACAAAGTCAATTCATAGTAGATCTTGGAGCTTACCTAGAAAAAGAAAGTATATATT
ATGTTACCACAAATTGGAAGCACACAGCTACACGATTTGCAATCAAACCCATGGAAGATTAACCTTTTTCCTCTACATCAGTGTGTTAATTCATACAAACTTTCTACCTACA
TTCTTCACTTCACCATCACAATCACAAACACTCTGTGGTTCAACCAATCAAACAAAACTTATCTGAAGTCCCAGATCATCCCAAGTCATTGTTTATCAGATCTAGTACTCAA
ATAAGTTAATAAAAAATATACACATGGGGCAAATAATCATTGGAGGAAATCCAACTAATCACAATATCTGTTAACATAGACAAGTCCACACACCATACAGAATCAACCAATG
GAAAATACATCCATAACAATAGAATTCTCAAGCAAATTCTGGCCTTACTTTACACTAATACACATGATCACAACAATAATCTCTTTGCTAATCATAATCTCCATCATGATTG
CAATACTAAACAAACTTTGTGAATATAACGTATTCCATAACAAAACCTTTGAGTTACCAAGAGCTCGAGTCAACACATAGCATTCATCAATCCAACAGCCCAAAACAGTAAC
CTTGCATTTAAAAATGAACAACCCCTACCTCTTTACAACACCTCATTAACATCCCACCATGCAAACCACTATCCATACTATAAAGTAGTTAATTAAAAATAGTCATAACAAT
GAACTAGGATATCAAGACTAACAATAACATTGGGGCAAATGCAAACATGTCCAAAAACAAGGACCAACGCACCGCTAAGACATTAGAAAGGACCTGGGACACTCTCAATCAT
TTATTATTCATATCATCGTGCTTATATAAGTTAAATCTTAAATCTGTAGCACAAATCACATTATCCATTCTGGCAATGATAATCTCAACTTCACTTATAATTGCAGCCATCA
TATTCATAGCCTCGGCAAACCACAAAGTCACACCAACAACTGCAATCATACAAGATGCAACAAGCCAGATCAAGAACACAACCCCAACATACCTCACCCAGAATCCTCAGCT
TGGAATCAGTCCCTCTAATCCGTCTGAAATTACATCACAAATCACCACCATACTAGCTTCAACAACACCAGGAGTCAAGTCAACCCTGCAATCCACAACAGTCAAGACCAAA
AACACAACAACAACTCAAACACAACCCAGCAAGCCCACCACAAAACAACGCCAAAACAAACCACCAAGCAAACCCAATAATGATTTTCACTTTGAAGTGTTCAACTTTGTAC
CCTGCAGCATATGCAGCAACAATCCAACCTGCTGGGCTATCTGCAAAAGAATACCAAACAAAAAACCAGGAAAGAAAACCACTACCAAGCCCACAAAAAAACCAACCCTCAA
GACAACCAAAAAAGATCCCAAACCTCAAACCACTAAATCAAAGGAAGTACCCACCACCAAGCCCACAGAAGAGCCAACCATCAACACCACCAAAACAAACATCATAACTACA
CTACTCACCTCCAACACCACAGGAAATCCAGAACTCACAAGTCAAATGGAAACCTTCCACTCAACTTCCTCCGAAGGCAATCCAAGCCCTTCTCAAGTCTCTACAACATCCG
AGTACCCATCACAACCTTCATCTCCACCCAACACACCACGCCAG**GGCGGCGGCGATTCTCTTGAATTTATTGCTAGTAAGCTTGCG**TAGTTACTTAAAAACATATTATCACA
AAAAGCCATGACCAACTTAAACAGAATCAAAATAAACTCTGGGGCAAATAACAATGGAGTTGCCAATCCTCAAAGCAAATGCAATTACCACAATCCTCGCTGCAGTCACATT
TTGCTTTGCTTCTGGT**GGCGGAGGAGGGGGC**CAAAACATCACTGAAGAATTTTATCAATCAACATGCAGTGCAGTTAGCAAAGGCTATCTTAGTGCTCTAAGAACTGGTTGG
TATACTAGTGTTATAACTATAGAATTAAGTAATATCAAGAAAAATAAGTGTAATGGAACAGATGCTAAGGTAAAATTGATGAAACAAGAATTAGATAAATATAAAAATGCTG
TAACAGAATTGCAGTTGCTCATGCAAAGCACACCAGCAGCAAACAATCGAGCCAGAAGAGAACTACCAAGGTTTATGAATTATACACTCAACAATACCAAAAAAACCAATGT
AACATTAAGCAAGAAAAGGAAAAGAAGATTTCTTGGTTTTTTGTTAGGTGTTGGATCTGCAATCGCCAGTGGCATTGCTGTATCTAAGGTCCTGCACTTAGAAGGAGAAGTG
AACAAGATCAAAAGTGCTCTACTATCCACAAACAAGGCCGTAGTCAGCTTATCAAATGGAGTTAGTGTCTTAACCAGCAGAGTGTTAGACCTCAAAAACTATATAGATAAAC
AATTGTTACCTATTGTGAATAAGCAAAGCTGCAGAATATCAAATATAGAAACTGTGATAGAGTTCCAACAAAAGAACAACAGACTACTAGAGATTACCAGGGAATTTAGTGT
TAATGCAGGTGTAACTACACCTGTAAGCACTTACATGTTAACTAATAGTGAATTATTGTCATTAATCAATGATATGCCTATAACAAATGATCAGAAAAAGTTAATGTCCAAC
AATGTTCAAATAGTTAGACAGCAAAGTTACTCTATCATGTCCATAATAAAAGAGGAAGTCTTAGCATATGTAGTACAATTACCACTATATGGTGTGATAGATACACCTTGTT
GGAAATTACACACATCCCCTCTATGTACAACCAACACAAAAGAAGGGTCAAACATCTGTTTAACAAGAACTGACAGAGGATGGTACTGTGACAATGCAGGATCAGTATCTTT
CTTCCCACAAGCTGAAAATGTAAAGTTCAATCGAATCGAGTATTTTGTGACACAATGTACAGTTTAACATTACCAAGTGAAGTAAATCTCTGCAATGTTGACATATTCAAT

CCCAAATATGATTGTAAAATTATGACTTCAAAAACAGATGTAAGCAGCTCCGTTATCACATCTCTAGGAGCCATTGTGTCATGCTATGGCAAAACTAAATGTACAGCATCCA
ATAAAAATCGTGGAATCATAAAGACATTTTCTAACGGGTGTGATTATGTATCAAATAAAGGGGTGGACACTGTGTCTGTAGGTAACACATTATATTATGTAAATAAGCAAGA
AGGCAAAAGTCTCTATGTAAAAGGTGAACCAATAATAAATTTCTATGACCCATTAGTATTCCCCTCTGATGAATTTGATGCATCAATATCTCAAGTCAATGAGAAGATTAAC
CAGAGTTTAGCATTTATTCGTAAATCCGATGAATTATTACATAATGTAAATGCTGGTAAATCAACCACAAATATCATGATAACTACTATAATTATAGTGATTATAGTAATAT
TGTTATCATTAATTGCTGTTGGACTGCTCCTATACTGTAAGGCCAGAAGCACACCAATCACACTAAGCAAGGATCAACTGAGTGGTATAAATAATATTGCATTTAGTAACTG
AATAAAAATAGCACCTAATCATGTTCTTACAATGGTTTACTATCTGCTCATAGACAACCCATCTATCATTGGATTTTCTTAAAATCTGAACTTCATCGAAACTCTTATCTAT
AAACCATCTCACTTACACTATTTAAGTAGATTCCTAGTTTATAGTTATATAAAACACAATTGAATGCCAGATTAACTTACCATCTGTAAAAATGAAAACTGGGGCAAATATG
TCACGAAGGAATCCTTGCAAATTTGAAATTCGAGGTCATTGCTTAAATGGTAAGAGGTGTCATTTTAGTCATAATTATTTTGAATGGCCACCCCATGCACTGCTTGTAAGAC
AAAACTTTATGTTAAACAGAATACTTAAGTCTATGGATAAAAGTATAGATACCTTATCAGAAATAAGTGGAGCTGCAGAGTTGGACAGAACAGAAGAGTATGCTCTTGGTGT
AGTTGGAGTGCTAGAGAGTTATATAGGATCAATAAACAATATAACTAAACAATCAGCATGTGTTGCCATGAGCAAACTCCTCACTGAACTCAATAGTGATGATATCAAAAAG
CTGAGGGACAATGAAGAGCTAAATTCACCCAAGATAAGAGTGTACAATACTGTCATATCATATATTGAAAGCAACAGGAAAAACAATAAACAAACTATCCATCTGTTAAAAA
GATTGCCAGCAGACGTATTGAAGAAAACCATCAAAAACACATTGGATATCCATAAGAGCATAACCATCAACAACCCAAAAGAATCAACTGTTAGTGATACAAATGACCATGC
CAAAAATAATGATACTACCTGACAAATATCCTTGTAGTATAACTTCCATACTAATAACAAGTAGATGTAGAGTTACTATGTATAATCAAAAGAACACACTATATTTCAATCA
AAACAACCCAAATAACCATATGTACTCACCGAATCAAACATTCAATGAAATCCATTGGACCTCTCAAGAATTGATTGACACAATTCAAATTTTTCTACAACATCTAGGTATT
ATTGAGGATATATATACAATATATATATTAGTGTCATAACACTCAATTCTAACACTCACCACATCGTTACATTATTAATTCAAACAATTCAAGTTGTGGGACAAAATGGATC
CCATTATTAATGGAAATTCTGCTAATGTTTATCTAACCGATAGTTATTTAAAAGGTGTTATCTCTTTCTCAGAGTGTAATGCTTTAGGAAGTTACATATTCAATGGTCCTTA
TCTCAAAAATGATTATACCAACTTAATTAGTAGACAAAATCCATTAATAGAACACATGAATCTAAAGAAACTAAATATAACACAGTCCTTAATATCTAAGTATCATAAAGGT
GAAATAAAATTAGAAGAACCTACTTATTTTCAGTCATTACTTATGACATACAAGAGTATGACCTCGTCAGAACAGATTGCTACCACTAATTTACTTAAAAAGATAATAAGAA
GAGCTATAGAAATAAGTGATGTCAAAGTCTATGCTATATTGAATAAACTAGGGCTTAAAGAAAAGGACAAGATTAAATCCAACAATGGACAAGATGAAGACAACTCAGTTAT
TACGACCATAATCAAAGATGATATACTTTCAGCTGTTAAAGATAATCAATCTCATCTTAAAGCAGACAAAAATCACTCTACAAAACAAAAAGACACAATCAAAACAACACTC
TTGAAGAAATTGATGTGTTCAATGCAACATCCTCCATCATGGTTAATACATTGGTTTAACTTATACACAAAATTAAACAACATATTAACACAGTATCGATCAAATGAGGTAA
AAAACCATGGGTTTACATTGATAGATAATCAAACTCTTAGTGGATTTCAATTTATTTTGAACCAATATGGTTGTATAGTTTATCATAAGGAACTCAAAAGAATTACTGTGAC
AACCTATAATCAATTCTTGACATGGAAAGATATTAGCCTTAGTAGATTAAATGTTTGTTTAATTACATGGATTAGTAACTGCTTGAACACATTAAATAAAAGCTTAGGCTTA
AGATGCGGATTCAATAATGTTATCTTGACACAACTATTCCTTTATGGAGATTGTATACTAAAGCTATTTCACAATGAGGGGTTCTACATAATAAAAGAGGTAGAGGGATTTA
TTATGTCTCTAATTTTAAATATAACAGAAGAAGATCAATTCAGAAAACGATTTTATAATAGTATGCTCAACAACATCACAGATGCTGCTAATAAAGCTCAGAAAAATCTGCT
ATCAAGAGTATGTCATACATTATTAGATAAGACAGTGTCCGATAATATAATAAATGGCAGATGGATAATTCTATTAAGTAAGTTCCTTAAATTAATTAAGCTTGCAGGTGAC
AATAACCTTAACAATCTGAGTGAACTATATTTTTTGTTCAGAATATTTGGACACCCAATGGTAGATGAAAGACAAGCCATGGATGCTGTTAAAATTAATTGCAATGAGACCA
AATTTTACTTGTTAAGCAGTCTGAGTATGTTAAGAGGTGCCTTTATATATAGAATTATAAAAGGGTTTGTAAATAATTACAACAGATGGCCTACTTTAAGAAATGCTATTGT
TTTACCCTTAAGATGGTTAACTTACTATAAACTAAACACTTATCCTTCTTTGTTGGAACTTACAGAAAGAGATTTGATTGTGTTATCAGGACTACGTTTCTATCGTGAGTTT
CGGTTGCCTAAAAAAGTGGATCTTGAAATGATTATAAATGATAAAGCTATATCACCTCCTAAAAATTTGATATGGACTAGTTTCCCTAGAAATTACATGCCATCACACATAC
AAAACTATATAGAACATGAAAAATTAAAATTTTCCGAGAGTGATAAATCAAGAAGAGTATTAGAGTATTATTTAAGAGATAACAAATTCAATGAATGTGATTTATACAACTG
TGTAGTTAATCAAAGTTATCTCAACAACCCTAATCATGTGGTATCATTGACAGGCAAAGAAAGAGAACTCAGTGTAGGTAGAATGTTTGCAATGCAACCGGGAATGTTCAGA
CAGGTTCAAATATTGGCAGAGAAAATGATAGCTGAAAACATTTTACAATTCTTTCCTGAAAGTCTTACAAGATATGGTGATCTAGAACTACAAAAAATATTAGAATTGAAAG
CAGGAATAAGTAACAAATCAAATCGCTACAATGATAATTACAACAATTACATTAGTAAGTGCTCTATCATCACAGATCTCAGCAAATTCAATCAAGCATTTCGATATGAAAC
GTCATGTATTTGTAGTGATGTGCTGGATGAACTGCATGGTGTACAATCTCTATTTTCCTGGTTACATTTAACTATTCCTCATGTCACAATAATATGCACATATAGGCATGCA
CCCCCCTATATAGGAGATCATATTGTAGATCTTAACAATGTAGATGAACAAAGTGGATTATATAGATATCACATGGGTGGCATCGAAGGGTGGTGTCAAAAACTGTGGACCA
TAGAAGCTATATCACTATTGGATCTAATATCTCTCAAAGGGAAATTCTCAATTACTGCTTTAATTAATGGTGACAATCAATCAATAGATATAAGCAAACCAATCAGACTCAT
GGAAGGTCAAACTCATGCTCAAGCAGATTATTTGCTAGCATTAAATAGCCTTAAATTACTGTATAAAGAGTATGCAGGCATAGGCCACAAATTAAAAGGAACTGAGACTTAT
ATATCACGAGATATGCAATTTATGAGTAAAACAATTCAACATAACGGTGTATATTACCCAGCTAGTATAAAGAAAGTCCTAAGAGTGGGACCGTGGATAAACACTATACTTG
ATGATTTCAAAGTGAGTCTAGAATCTATAGGTAGTTTGACACAAGAATTAGAATATAGAGGTGAAAGTCTATTATGCAGTTTAATATTTAGAAATGTATGGTTATATAATCA
GATTGCTCTACAATTAAAAAATCATGCATTATGTAACAATAAACTATATTTGGACATATTAAAGGTTCTGAAACACTTAAAAACCTTTTTTAATCTTGATAATATTGATACA
GCATTAACATTGTATATGAATTTACCCATGTTATTTGGTGGTGGTGATCCCAACTTGTTATATCGAAGTTTCTATAGAAGAACTCCTGACTTCCTCACAGAGGCTATAGTTC
ACTCTGTGTTCATACTTAGTTATTATACAAACCATGACTTAAAAGATAAACTTCAAGATCTGTCAGATGATAGATTGAATAAGTTCTTAACATGCATAATCACGTTTGACAA
AAACCCTAATGCTGAATTCGTAACATTGATGAGAGATCCTCAAGCTTTAGGGTCTGAGAGACAAGCTAAAATTACTAGCGAAATCAATAGACTGGCAGTTACAGAGGTTTTG
AGTACAGCTCCAAACAAAATATTCTCCAAAAGTGCACAACATTATACTACTACAGAGATAGATCTAAATGATATTATGCAAAATATAGAACCTACATATCCTCATGGGCTAA
GAGTTGTTTATGAAAGTTTACCCTTTTATAAAGCAGAGAAAATAGTAAATCTTATATCAGGTACAAAATCTATAACTAACATACTGGAAAAAACTTCTGCCATAGACTTAAC
AGATATTGATAGAGCCACTGAGATGATGAGGAAAAACATAACTTTGCTTATAAGGATACTTCCATTGGATTGTAACAGAGATAAAAGAGAGATATTGAGTATGGAAAACCTA
AGTATTACTGAATTAAGCAAATATGTTAGGGAAAGATCTTGGTCTTTATCCAATATAGTTGGTGTTACATCACCCAGTATCATGTATACAATGGACATCAAATATACTACAA
GCACTATATCTAGTGGCATAATTATAGAGAAATATAATGTTAACAGTTTAACACGTGGTGAGAGAGGACCCACTAAACCATGGGTTGGTTCATCTACACAAGAGAAAAAAAC
AATGCCAGTTTATAATAGACAAGTCTTAACCAAAAAACAGAGAGATCAAATAGATCTATTAGCAAAATTGGATTGGGTGTATGCATCTATAGATAACAAGGATGAATTCATG
GAAGAACTCAGCATAGGAACCCTTGGGTTAACATATGAAAAGGCCAAGAAATTATTTCCACAATATTTAAGTGTCAATTATTTGCATCGCCTTACAGTCAGTAGTAGACCAT
GTGAATTCCCTGCATCAATACCAGCTTATAGAACAACAAATTATCACTTTGACACTAGCCCTATTAATCGCATATTAACAGAAAAGTATGGTGATGAAGATATTGACATAGT
ATTCCAAAACTGTATAAGCTTTGGCCTTAGTTTAATGTCAGTAGTAGAACAATTTACTAATGTATGTCCTAACAGAATTATTCTCATACCTAAGCTTAATGAGATACATTTG
ATGAAACCTCCCATATTCACAGGTGATGTTGATATTCACAAGTTAAAACAAGTGATACAAAAACAGCATATGTTTTTACCAGACAAAATAAGTTTGACTCAATATGTGGAAT
TATTCTTAAGTAATAAAACACTCAAATCTGGATCTCATGTTAATTCTAATTTAATATTGGCACATAAAATATCTGACTATTTTCATAATACTTACATTTTAAGTACTAATTT
AGCTGGACATTGGATTCTGATTATACAACTTATGAAAGATTCTAAAGGTATTTTTGAAAAAGATTGGGGAGAGGGATATATAACTGATCATATGTTTATTAATTTGAAAGTT
TTCTTCAATGCTTATAAGACCTATCTCTTGTGTTTTCATAAAGGTTATGGCAAAGCAAAGCTGGAGTGTGATATGAACACTTCAGATCTTCTATGTGTATTGGAATTAATAG
ACAGTAGTTATTGGAAGTCTATGTCTAAGGTATTTTTAGAACAAAAAGTTATCAAATACATTCTTAGCCAAGATGCAAGTTTACATAGAGTAAAAGGATGTCATAGCTTCAA
ATTATGGTTTCTTAAACGTCTTAATGTAGCAGAATTCACAGTTTGCCCTTGGGTTGTTAACATAGATTATCATCCAACACATATGAAAGCAATATTAACTTATATAGATCTT
GTTAGAATGGGATTGATAAATATAGATAGAATACACATTAAAAATAAACACAAATTCAATGATGAATTTTATACTTCTAATCTCTTCTACATTAATTATAACTTCTCAGATA
ATACTCATCTATTAACTAAACATATAAGGATTGCTAATTCTGAATTAGAAAATAATTACAACAAATTATATCATCCTACACCAGAAACCCTAGAGAATATACTAGCCAATCC
GATTAAAAGTAATGACAAAAAGACACTGAATGACTATTGTATAGGTAAAAATGTTGACTCAATAATGTTACCATTGTTATCTAATAAGAAGCTTATTAAATCGTCTGCAATG
ATTAGAACCAATTACAGCAAACAAGATTTGTATAATTTATTCCCTATGGTTGTGATTGATAGAATTATAGATCATTCAGGCAATACAGCCAAATCCAACCAACTTTACACTA
CTACTTCCCACCAAATATCTTTAGTGCACAATAGCACATCACTTTACTGCATGCTTCCTTGGCATCATATTAATAGATTCAATTTTGTATTTAGTTCTACAGGTTGTAAAAT
TAGTATAGAGTATATTTTAAAAGATCTTAAAATTAAAGATCCCAATTGTATAGCATTCATAGGTGAAGGAGCAGGGAATTTATTATTGCGTACAGTAGTGGAACTTCATCCT
GACATAAGATATATTTACAGAAGTCTGAAAGATTGCAATGATCATAGTTTACCTATTGAGTTTTTAAGGCTGTACAATGGACATATCAACATTGATTATGGTGAAAATTTGA
CCATTCCTGCTACAGATGCAACCAACAACATTCATTGGTCTTATTTACATATAAAGTTTGCTGAACCTATCAGTCTTTTTGTCTGTGATGCCGAATTGTCTGTAACAGTCAA
CTGGAGTAAAATTATAATAGAATGGAGCAAGCATGTAAGAAAGTGCAAGTACTGTTCCTCAGTTAATAAATGTATGTTAATAGTAAAATATCATGCTCAAGATGATATTGAT
TTCAAATTAGACAATATAACTATATTAAAAACTTATGTATGCTTAGGCAGTAAGTTAAAGGGATCGGAGGTTTACTTAGTCCTTACAATAGGTCCTGCGAATATATTCCCAG
TATTTAATGTAGTACAAAATGCTAAATTGATACTATCAAGAACCAAAAATTTCATCATGCCTAAGAAAGCTGATAAAGAGTCTATTGATGCAAATATTAAAAGTTTGATACC
CTTTCTTTGTTACCCTATAACAAAAAAAGGAATTAATACTGCATTGTCAAAACTAAAGAGTGTTGTTAGTGGAGATATACTATCATATTCTATAGCTGGACGTAATGAAGTT
TTCAGCAATAAACTTATAAATCATAAGCATATGAACATCTTAAAATGGTTCAATCATGTTTTAAATTTCAGATCAACAGAACTAAACTATAACCATTTATATATGGTAGAAT
CTACATATCCTTACCTAAGTGAATTGTTAAACAGCTTGACAACCAATGAACTTAAAAAACTGATTAAAATCACAGGTAGTCTGTTATACAACTTTCATAATGAATAATGAAT
AAAGATCTTATAATAAAAATTCCCATAGCTATACACTAACACTGTATTCAATTATAGTTATTAAAAATTAAAAATCATATAATTTTTTAAATAACTTTTAGTGAACTAATCC
TAAAGTTATCATTTTAATCTTGGAGGAATAAATTTAAACCCTAATCTAATTGGTTTATATGTGTATTAACTAAATTACGAGATATTAGTTTTTGACACTTTTTTTCTCGT

### Antibody sequences

Signal sequence / VH/VL / CH/CL / linker / ybbR-tag / affinity tag

CR9501 HC:

MGWSCIILFLVATATGVHSQVQLVQSGPGLVKPSQTLALTCTVSGASINSDNYYWTWIRQRPGGGLEWIGHISYTGNTYYTPSLKSRLSMSLETSQSQFSLRLTSVTAADSAVYFCAACGAYVLISNCGWFDSWGQGTQVTVSSASTKGPSVFPLAPSSKSTSGGTAALGCLVKDYFPEPVTVSWNSGALTSGVHTFPAVLQSSGLYSLSSVVTVPSSSLGTQTYICNVNHKPSNTKVDKRVEPKSCDKTHTCPPCPAPELLGGPSVFLFPPKPKDTLMISRTPEVTCVVVDVSHEDPEVKFNWYVDGVEVHNAKTKPREEQYNSTYRVVSVLTVLHQDWLNGKEYKCKVSNKALPAPIEKTISKAKGQPREPQVYTLPPSRDELTKNQVSLTCLVKGFYPSDIAVEWESNGQPENNYKTTPPVLDSDGSFFLYSKLTVDKSRWQQGNVFSCSVMHEALHNHYTQKSLSLSPGKGGSGGSDSLEFIASKLA*

CR9501 LC:

MGWSCIILFLVATATGVHSEIVMTQSPSSLSASIGDRVTITCQASQDISTYLNWYQQKPGQAPRLLIYGASNLETGVPSRFTGSGYGTDFSVTISSLQPEDIATYYCQQYQYLPYTFAPGTKVEIKRTVAAPSVFIFPPSDEQLKSGTASVVCLLNNFYPREAKVQWKVDNALQSGNSQESVTEQDSKDSTYSLSSTLTLSKADYEKHKVYACEVTHQGLSSPVTKSFNRGEC

5C4 HC:

MGWSCIILFLVATATGVHSVQLQQSGAELVKPGASVKLSCTASGFNIKDTFFHWVKQRPEQGLEWIGRIDPADGHTKYDPKFQGKATITADTSSNTAFLQLSSLTSVDTAVYYCATTITAVVPTPYNAMDYWGQGTTVTVSSASTKGPSVFPLAPSSKSTSGGTAALGCLVKDYFPEPVTVSWNSGALTSGVHTFPAVLQSSGLYSLSSVVTVPSSSLGTQTYICNVNHKPSNTKVDKRVEPKSCDKTHTCPPCPAPELLGGPSVFLFPPKPKDTLMISRTPEVTCVVVDVSHEDPEVKFNWYVDGVEVHNAKTKPREEQYNSTYRVVSVLTVLHQDWLNGKEYKCKVSNKALPAPIEKTISKAKGQPREPQVYTLPPSRDELTKNQVSLTCLVKGFYPSDIAVEWESNGQPENNYKTTPPVLDSDGSFFLYSKLTVDKSRWQQGNVFSCSVMHEALHNHYTQKSLSLSPGKGGSGGSDSLEFIASKLA*

5C4 LC:

MGWSCIILFLVATATGVHSDIVLTQSPASLAVSLGQRTTISCRASESVDSFDNSFIHWYQQKPGQPPKLLIFLASSLESGVPARFSGSGSRTDFTLTIDPVEADDAATYYCQQSNEDPFTFGSGTKLEIKRADAAPSVFIFPPSDEQLKSGTASVVCLLNNFYPREAKVQWKVDNALQSGNSQESVTEQDSKDSTYSLSSTLTLSKADYEKHKVYACEVTHQGLSSPVTKSFNRGEC*

Motavizumab HC:

MGWSCIILFLVATATGVHSQVTLRESGPALVKPTQTLTLTCTFSGFSLSTAGMSVGWIRQPPGKALEWLADIWWDDKKHYNPSLKDRLTISKDTSKNQVVLKVTNMDPADTATYYCARDMIFNFYFDVWGQGTTVTVSSASTKGPSVFPLAPSSKSTSGGTAALGCLVKDYFPEPVTVSWNSGALTSGVHTFPAVLQSSGLYSLSSVVTVPSSSLGTQTYICNVNHKPSNTKVDKRVEPKSCDKTHTCPPCPAPELLGGPSVFLFPPKPKDTLMISRTPEVTCVVVDVSHEDPEVKFNWYVDGVEVHNAKTKPREEQYNSTYRVVSVLTVLHQDWLNGKEYKCKVSNKALPAPIEKTISKAKGQPREPQVYTLPPSRDELTKNQVSLTCLVKGFYPSDIAVEWESNGQPENNYKTTPPVLDSDGSFFLYSKLTVDKSRWQQGNVFSCSVMHEALHNHYTQKSLSLSPGKGGSGGSDSLEFIASKLA*

Motavizumab LC:

MGWSCIILFLVATATGVHSDIQMTQSPSTLSASVGDRVTITCSASSRVGYMHWYQQKPGKAPKLLIYDTSKLASGVPSRFSGSGSGTEFTLTISSLQPDDFATYYCFQGSGYPFTFGGGTKVEIKRTVAAPSVFIFPPSDEQLKSGTASVVCLLNNFYPREAKVQWKVDNALQSGNSQESVTEQDSKDSTYSLSSTLTLSKADYEKHKVYACEVTHQGLSSPVTKSFNRGEC*

101F HC:

MGWSCIILFLVATATGVHSQVTLKESGPGILQPSQTLSLTCSFSGFSLSTSGMGVSWIRQPSGKGLEWLAHIYWDDDKRYNPSLKSRLTISKDTSRNQVFLKITSVDTADTATYYCARLYGFTYGFAYWGQGTLVTVSSASTKGPSVFPLAPSSKSTSGGTAALGCLVKDYFPEPVTVSWNSGALTSGVHTFPAVLQSSGLYSLSSVVTVPSSSLGTQTYICNVNHKPSNTKVDKRVEPKSCDKTHTCPPCPAPELLGGPSVFLFPPKPKDTLMISRTPEVTCVVVDVSHEDPEVKFNWYVDGVEVHNAKTKPREEQYNSTYRVVSVLTVLHQDWLNGKEYKCKVSNKALPAPIEKTISKAKGQPREPQVYTLPPSRDELTKNQVSLTCLVKGFYPSDIAVEWESNGQPENNYKTTPPVLDSDGSFFLYSKLTVDKSRWQQGNVFSCSVMHEALHNHYTQKSLSLSPGKGGSGGSDSLEFIASKLA*

101F LC:

MGWSCIILFLVATATGVHSDIVLTQSPASLAVSLGQRATIFCRASQSVDYNGISYMHWFQQKPGQPPKLLIYAASNPESGIPARFTGSGSGTDFTLNIHPVEEEDAATYYCQQIIEDPWTFGGGTKLEIKRADAAPSVFIFPPSDEQLKSGTASVVCLLNNFYPREAKVQWKVDNALQSGNSQESVTEQDSKDSTYSLSSTLTLSKADYEKHKVYACEVTHQGLSSPVTKSFNRGEC*

ADI-19425 HC:

MGWSCIILFLVATATGVHSEVQLVESGGGLVKPGGSLRLSCAASGFTFSSYSMNWVRQAPGKGLEWVSSISSSSSYIYYADSVKGRFTISRDNAKNSLYLQMNSLRAEDTAVYYCARLGYCSGGSCHFDYWGQGTLVTVSSASTKGPSVFPLAPSSKSTSGGTAALGCLVKDYFPEPVTVSWNSGALTSGVHTFPAVLQSSGLYSLSSVVTVPSSSLGTQTYICNVNHKPSNTKVDKRVEPKSCDKTHTCPPCPAPELLGGPSVFLFPPKPKDTLMISRTPEVTCVVVDVSHEDPEVKFNWYVDGVEVHNAKTKPREEQYNSTYRVVSVLTVLHQDWLNGKEYKCKVSNKALPAPIEKTISKAKGQPREPQVYTLPPSRDELTKNQVSLTCLVKGFYPSDIAVEWESNGQPENNYKTTPPVLDSDGSFFLYSKLTVDKSRWQQGNVFSCSVMHEALHNHYTQKSLSLSPGKGGSGGSDSLEFIASKLA*

ADI-19425 LC:

MGWSCIILFLVATATGVHSQPVLTQPPSVSGAPGQRVTISCTGSSSNIGAGYDVHWYQQLPGTAPKLLIYGNSNRPSGVPDRFSGSKSGTSASLAITGLQAEDEADYYCQSYDSSLSGFYVFGTGTKLTVLGQPKAAPSVFIFPPSDEQLKSGTASVVCLLNNFYPREAKVQWKVDNALQSGNSQESVTEQDSKDSTYSLSSTLTLSKADYEKHKVYACEVTHQGLSSPVTKSFNRGEC*

ADI-14353 HC:
MGWSCIILFLVATATGVHSQVQLVQSGAEVKKPGSSVKVSCKASGGTFSSYTISWVRQAPGQGLEWMGRIKPIIGIANNAQRFKGRVTI
TAEKSTGTAYMELSSLTSEDTAVYYCARGGYDYYGMDVWGQGTTVTVSSASTKGPSVFPLAPSSKSTSGGTAALGCLVKDYFPEPVTVS
WNSGALTSGVHTFPAVLQSSGLYSLSSVVTVPSSSLGTQTYICNVNHKPSNTKVDKRVEPKSCDKTHTCPPCPAPELLGGPSVFLFPPK
PKDTLMISRTPEVTCVVVDVSHEDPEVKFNWYVDGVEVHNAKTKPREEQYNSTYRVVSVLTVLHQDWLNGKEYKCKVSNKALPAPIEKT
ISKAKGQPREPQVYTLPPSRDELTKNQVSLTCLVKGFYPSDIAVEWESNGQPENNYKTTPPVLDSDGSFFLYSKLTVDKSRWQQGNVFS
CSVMHEALHNHYTQKSLSLSPGKGGSGGSDSLEFIASKLA*
ADI-14353 LC:
MGWSCIILFLVATATGVHSQSALTQPASVSGSPGQSITISCTGTSSDVGGYNYVSWHQQHPGKAPKLLIYDVSNRPSGVSNRFSGSKSG
NTASLSISGLQAEDEADYYCSSFTSTSTPYVFGTGTQLTVLGQPKAAPSVFIFPPSDEQLKSGTASVVCLLNNFYPREAKVQWKVDNAL
QSGNSQESVTEQDSKDSTYSLSSTLTLSKADYEKHKVYACEVTHQGLSSPVTKSFNRGEC*

ADI-14359 HC:
MGWSCIILFLVATATGVHSQVTLRESGPALVKPTQTLTLTCTFSGFSLSTSGMCVSWIRQPPGKALEWLARIDWDDDKYYSTSLKTRLT
ISKDTSKNQVVLTMTNMDPVDTATYYCARATNYDSSGYYSLYFDYWGQGTLVTVSSASTKGPSVFPLAPSSKSTSGGTAALGCLVKDYF
PEPVTVSWNSGALTSGVHTFPAVLQSSGLYSLSSVVTVPSSSLGTQTYICNVNHKPSNTKVDKRVEPKSCDKTHTCPPCPAPELLGGPS
VFLFPPKPKDTLMISRTPEVTCVVVDVSHEDPEVKFNWYVDGVEVHNAKTKPREEQYNSTYRVVSVLTVLHQDWLNGKEYKCKVSNKAL
PAPIEKTISKAKGQPREPQVYTLPPSRDELTKNQVSLTCLVKGFYPSDIAVEWESNGQPENNYKTTPPVLDSDGSFFLYSKLTVDKSRW
QQGNVFSCSVMHEALHNHYTQKSLSLSPGKGGSGGSDSLEFIASKLA*
ADI-14359 LC:
MGWSCIILFLVATATGVHSDIQMTQSPSSLSASVGDRVTITCRASQSISSYLNWYQQKPGKAPKLLIYAASSLQSGVPSRFSGSGSGTD
FTLTISSLQPEDFATYYCQQSYSTPYTFGGGTKVEIKRTVAAPSVFIFPPSDEQLKSGTASVVCLLNNFYPREAKVQWKVDNALQSGNS
QESVTEQDSKDSTYSLSSTLTLSKADYEKHKVYACEVTHQGLSSPVTKSFNRGEC*

3D3 HC:
MGWSCIILFLVATATGVHSEEQLVESGGGLVQPGRSLRLSCVGSGLRFEEHAMHWVRQAPGRGLEWVSGISWNSGSVGYADSVKGRFTT
SRDNAKDILFLEMNTLRSEDTALYFCAIMVATTKNDFHYYKDVWGKGTTVTVSSASTKGPSVFPLAPSSKSTSGGTAALGCLVKDYFPE
PVTVSWNSGALTSGVHTFPAVLQSSGLYSLSSVVTVPSSSLGTQTYICNVNHKPSNTKVDKRVEPKSCDKTHTCPPCPAPELLGGPSVF
LFPPKPKDTLMISRTPEVTCVVVDVSHEDPEVKFNWYVDGVEVHNAKTKPREEQYNSTYRVVSVLTVLHQDWLNGKEYKCKVSNKALPA
PIEKTISKAKGQPREPQVYTLPPSRDELTKNQVSLTCLVKGFYPSDIAVEWESNGQPENNYKTTPPVLDSDGSFFLYSKLTVDKSRWQQ
GNVFSCSVMHEALHNHYTQKSLSLSPGKGGSGGSDSLEFIASKLA*
3D3 LC:
MGWSCIILFLVATATGVHSQIVLTQSPATLSLSPGERATLSCRASQSVSNHLAWYQQKPGQAPRLLIYETSNRATGIPPRFSGSGSGTD
FTLTISSLEPEDFAVYYCQQRNNWYTFGQGTKLEIKRTVAAPSVFIFPPSDEQLKSGTASVVCLLNNFYPREAKVQWKVDNALQSGNSQ
ESVTEQDSKDSTYSLSSTLTLSKADYEKHKVYACEVTHQGLSSPVTKSFNRGEC*

5C4 IgM HC:
MGWSCIILFLVATATGVHSVQLQQSGAELVKPGASVKLSCTASGFNIKDTFFHWVKQRPEQGLEWIGRIDPADGHTKYDPKFQGKATIT
ADTSSNTAFLQLSSLTSVDTAVYYCATTITAVVPTPYNAMDYWGQGTTVTVSSASASAPTLFPLVSCENSPSDTSSVAVGCLAQDFLPD
SITFSWKYKNNSDISSTRGFPSVLRGGKYAATSQVLLPSKDVMQGTDEHVVCKVQHPNGNKEKNVPLPVIAELPPKVSVFVPPRDGFFG
NPRKSKLICQATGFSPRQIQVSWLREGKQVGSGVTTDQVQAEAKESGPTTYKVTSTLTIKESDWLGQSMFTCRVDHRGLTFQQNASSMC
VPDQDTAIRVFAIPPSFASIFLTKSTKLTCLVTDLTTYDSVTISWTRQNGEAVKTHTNISESHPNATFSAVGEASICEDDWNSGERFTC
TVTHTDLPSPLKQTISRPKGVALHRPDVYLLPPAREQLNLRESATITCLVTGFSPADVFVQWMQRGQPLSPEKYVTSAPMPEPQAPGRY
FAHSILTVSEEEWNTGETYTCVVAHEALPNRVTERTVDKSTGKPTLYNVSLVMSDTAGTCYGGSHHHHHHGGSDSLEFIASKLA*

J Chain:
MGWSCIILFLVATATGVHSQEDERIVLVDNKCKCARITSRIIRSSEDPNEDIVERNIRIIVPLNNRENISDPTSPLRTRFVYHLSDLCK
KCDPTEVELDNQIVTATQSNICDEDSATETCYTYDRNKCYTAVVPLVYGGETKMVETALTPDACYPDGGSSGSNWSHPQFEKGGGGSNW
SHPQFEK*

ADI-14359 Fab HC:
MGWSCIILFLVATATGVHSQVTLRESGPALVKPTQTLTLTCTFSGFSLSTSGMCVSWIRQPPGKALEWLARIDWDDDKYYSTSLKTRLT
ISKDTSKNQVVLTMTNMDPVDTATYYCARATNYDSSGYYSLYFDYWGQGTLVTVSSASTKGPSVFPLAPSSKSTSGGTAALGCLVKDYF
PEPVTVSWNSGALTSGVHTFPAVLQSSGLYSLSSVVTVPSSSLGTQTYICNVNHKPSNTKVDKRVEPKSCDKGGSHHHHHHGGSDSLEF
IASKLA*
